# Integrated genotyping, annotation and epigenetic characterization of transposable elements with GraffiTE

**DOI:** 10.64898/2026.11.06.764993

**Authors:** Clément Goubert, Rose Laflamme, Robert J.M. Eveleigh, Jiannis Ragoussis, Travis J. Wheeler, Guillaume Lettre, Guillaume Bourque, Cristian Groza

## Abstract

Genotyping, annotating and characterizing TE variation leaped forward with long-read sequencing (LRS) and *de novo* assembly. LRS techniques also reveal CpG methylation, which enables measuring the impact of TEs on epigenomes and sheds light on the role of TEs as regulatory sequences. Yet, TE genotyping tools discard the valuable allele-specific epigenetic information.

We introduce GraffiTE v1.1, the first Nextflow pipeline that genotypes SVs, annotates polymorphic mobile elements (MEI), and maps epigenetic data to TEs using pangenome graphs in one command. GraffiTE v1.1 is designed to map any epigenetic information that is based on LRS stencils, including Fiber-seq. We demonstrate the capabilities of GraffiTE v1.1 by analyzing genomes and methylomes obtained from 20 Quebec residents of French-Canadian descent.

In these genomes, TE annotation yields 4,908 Alu, 704 LINE1, 112 SVA and 21 HERVK (HML-2) distinct polymorphic mobile elements (MEIs). We also identify 10 fixed HERVK insertions that vary between solo LTR and provirus, or in proviral copy number. We determine that these MEIs contribute 144,748 additional CpGs not present in the reference. From this, we identify 480 Alus, 67 LINE1s, 12 SVAs and 2 HERVKs in hypomethylated states. We note that Alus and SVAs are more methylated than other LINE1 and HERVK. Lastly, MEIs dating reported a mean of age 1.08 Mya (median: 640 kya) and beta regression identified a moderate demethylating effect of age on MEIs (effect: -0.069 log-odds per log10 years, P < 6.08x10^-19^). Our results show that the methylation level of a LINE1 MEI is expected to decrease from 86.4% to 80.8% in the first Mya.

In conclusion, GraffiTE v1.1 genotypes SVs, annotates MEI, and characterizes their epigenetic state at the scale of populations by leveraging the full potential of LRS. While we showcase its application in human data, the pipeline can be deployed in any model for which a TE library is available.

## Introduction

Epigenetic modification of DNA appears to be universal in eukaryotic genomes. For example, the methylation of cytosine is used to regulate the activity of genes (Breiling and Lyko 2015; Dhar et al. 2021) and the methylation of wide regions of DNA is used to silence parts of the genome (Moore et al. 2013). Moreover, the epigenome is not limited to methylation and extends to many features such as histone modifications and chromatin accessibility (Chen et al. 2024; Daugherty et al. 2017; Inbar-Feigenberg et al. 2013).

Transposable elements (TEs) are another shared feature of eukaryotic genomes. These elements propagate through genomes and contribute an important proportion of their sequence. Incidentally, TEs are silenced through epigenetic mechanisms to limit their proliferation. Despite their selfish nature, some TEs evolved to become regulatory elements and show enrichment in active epigenetic marks (Su et al. 2014) and have been associated with gene expression (Groza et al. 2023; Goubert et al. 2020). Therefore, epigenetic characterization of transposable elements is increasingly pursued to understand the co-evolution of genomes and epigenomes (Daron and Slotkin 2017; Zhuo et al. 2026; Merenciano et al. 2024; Zhuo et al. 2020; Ferraj et al. 2023).

Moreover, the activity of TEs is one of the drivers of structural variants (SVs), particularly through insertion polymorphisms, which are also referred to as (segregating) mobile element insertion (MEI, Goubert et al. 2020). MEI genotyping has been challenging in short-read next-generation sequencing (NGS) data and typically requires specialized tools to infer their presence. Due to the limitations of NGS, the TE genotype was often obtained without its sequence (Gardner et al. 2017; Chu et al. 2021). Notably, these young polymorphic insertions have been surmised to impact the epigenome at the site of insertion (Zhuo et al. 2026). However, the repetitive nature of TEs impeded mapping epigenetic data, especially when the TE length greatly exceeds the read length (Taylor et al. 2022; Ewing et al. 2020).

The emergence of long-read sequencing (LRS) technologies simplified this by yielding their full sequence (Groza et al. 2024; Mohamed et al. 2023; Cuenca-Guardiola et al. 2025). Even better, LRS technologies such as ONT and PacBio simultaneously reveal nucleotide modifications. In fact, novel techniques based on long-read stencils (the enzymatic labeling of adenine in single molecules) can extend the advantages of LRS to profiling histone modifications (Yue et al. 2022) and chromatin accessibility (Stergachis et al. 2020). Thus, LRS approaches can map diverse epigenomic features within long TE sequences that were previously unmappable with NGS approaches such as ChIP-seq (Ewing et al. 2020). At the same time, LRS is increasingly being recognized as a platform for epigenetics in ecology and evolution (de Carvalho et al. 2026).

The same LRS developments enabled large sets of high quality *de novo* assemblies of human genomes (Liao et al. 2023), and of model (Helmy et al. 2026) and non-model organisms (Armstrong et al. 2020; Leonard et al. 2023). Shortly, the Human Pangenome Reference Consortium (HPRC), released hundreds of *de novo* assemblies, compiled into the first human pangenome, which immediately proved to be a valuable resource for the study of SVs and their epigenetic properties (Lucas et al. 2026). As a result, methods like methylGrapher (Zhang et al. 2025) and panmethyl (Groza et al. 2025) have been developed to characterize the epigenetic state of pangenomes.

However, the integration of TE genotyping and annotation with epigenetic techniques based on LRS is lacking. Currently, taking full advantage of LRS to routinely study the epigenetics of TEs requires a demanding skill set in LRS, genome assembly, SV calling, TE annotation and epigenetics that is not widely available in the research community. To accelerate research in this area, we introduce GraffiTE v1.1, a Nextflow (Di Tommaso et al. 2017) pipeline that genotypes SVs, annotates TEs, and maps epigenetic data to TEs using pangenome graphs (Eizenga et al. 2020). GraffiTE v1.1 integrates the above steps into one command and produces VCFs containing TE polymorphisms annotated with their epigenetic state. Moreover, GraffiTE v1.1 is agnostic to the nature of the epigenetic assays, and is designed to map any epigenetic information that is based on LRS stencils (Jha et al. 2023).

Here, we demonstrate these capabilities on a new data set of 20 French-Canadian genomes and methylomes that were sequenced with PacBio HiFi (Hon et al. 2020) using samples from the CARTaGENE cohort (McClelland et al. 2026). After leveraging the novel features of the pipeline to identify *bona fide* insertion polymorphisms (resulting from transposition), including complex alleles (e.g. LINE1 5’ inversions, HERVK solo-LTR vs proviral insertion), we build on the phased haplotypes of the CARTaGENE dataset to estimate MEI insertion dates averaging ∼1 Mya. Layering methylation data from graph-genotyping, we identify hypomethylated outliers loci while modeling ∼5% demethylation pattern at segregating LINE1 during the last million years.

## Results

### A reliable set of SVs from CARTaGENE long-read sequencing

To demonstrate the capabilities of GraffiTE in calling, annotating and mapping the epigenetic state of TEs, we sequenced 20 samples of French-Canadian ancestry from Quebec’s CARTaGENE cohort with PacBio HiFi at a minimum coverage of 30X (Fig 1A). Upon assembly, we obtained 20 diploid genomes with an average N50 of 43.2 Mbp (min: 12.3 Mbp; max: 81.0 Mbp). The contiguity of the CARTaGENE assemblies is sufficient to detect most TEs given that human MEIs do not exceed 10 kbp.

**Fig 1.**
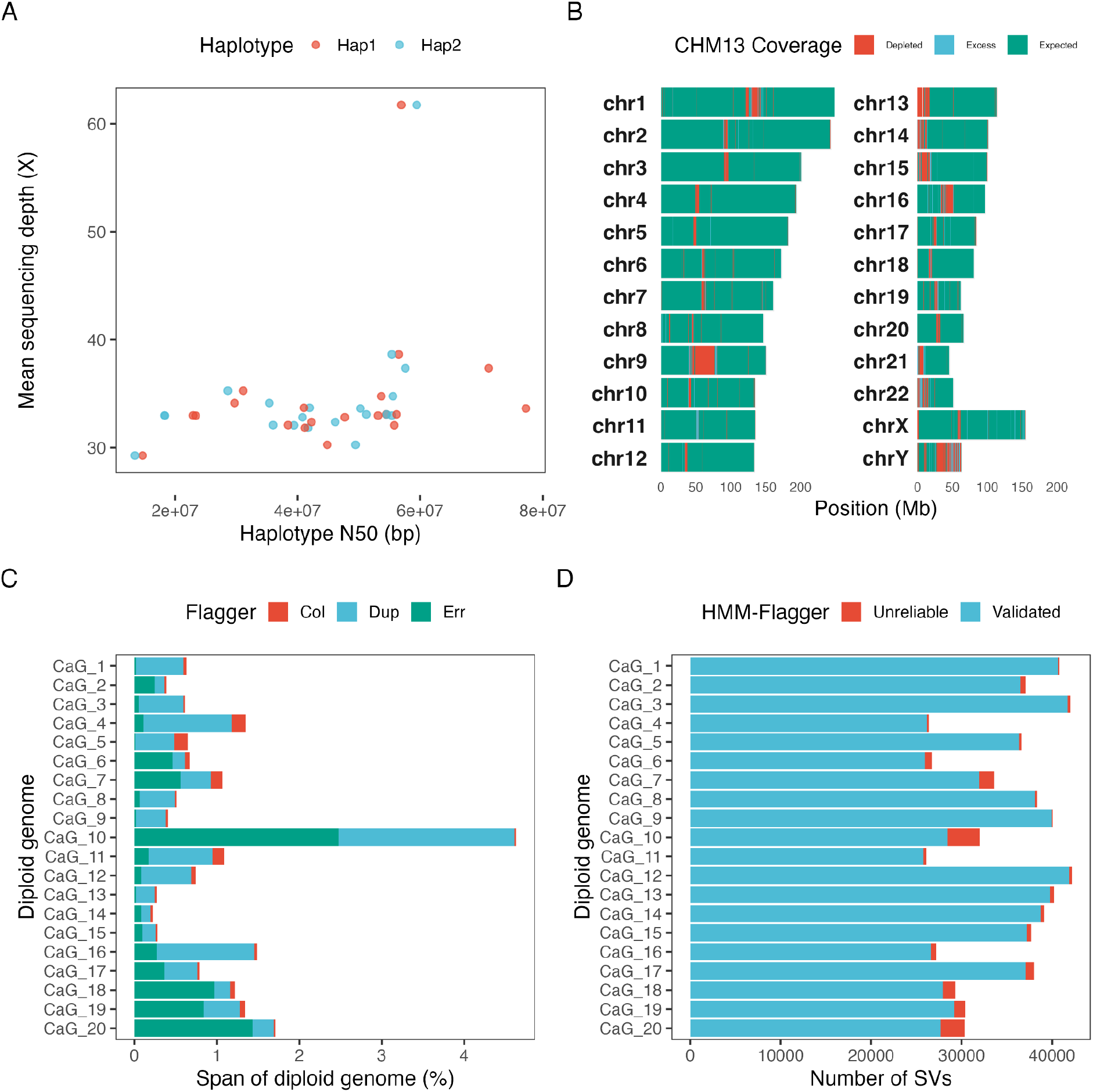
A) Sequencing depth and assembly contiguity of the CARTaGENE samples. B) Karyotype summarizing the alignments of CaG assemblies to CHM13. The expected coverage over the reference is 40 genomes. Regions with depleted coverage have been dropped in at least 10% of genomes. No assembly maps to white regions. CHM13 regions with excess coverage are 10% over the expectation. C) Validation with HMM-Flagger of the same assemblies, showing the unreliable portion of each assembly. D) Number of SVs detected with PAV. SVs supported by reliable and unreliable regions are highlighted.

To check the completeness of these diploid genomes, we aligned them against CHM13 (Fig 1B). We found that most regions are covered by all assemblies (Fig 1B). As expected, acrocentric arms, centromeres and the satellite on chromosome 9 are difficult to resolve with HiFi alone. Each diploid genome covered the vast majority of CHM13 (mean: 96.5%; min: 93.8%; max: 97.6%), indicating that we should recover most TE polymorphisms (Fig S1). Then, for the purposes of validation, we partitioned the assemblies with HMM-Flagger into reliable and unreliable sequences (Fig 1C). We found that the CARTaGENE assemblies are reliable, with only a small part of their span being flagged as being collapsed, duplicated or erroneous (mean: 1%; min: 0.22%; max: 4.63%).

To prepare an SV set for TE annotation, we called variants from each diploid assembly against CHM13 (Methods). Prior to filtering, we counted 34,717 SVs per diploid genome (min: 26,072; max: 42,202). After excluding SVs supported by unreliable assembly sequences, we retained 33,886 SVs per diploid genome (min: 25,738; max: 41,943) and removed 831 putatively false positive SVs per diploid genome (min: 106; max: 3,585). We consolidated SVs across the 20 diploid genomes into a final set of 147,939 reliable SVs for TE annotation and epigenetic mapping. These results show that the GraffiTE workflow is transparent and allows integration with complementary methods to identify a set of reliable SVs.

### Comprehensive annotation of MEIs within the CARTaGENE cohort

To look for high-quality polymorphic mobile element variants (MEIs) from this list of SVs, we used the *--human* mode of GraffiTE1.1 which applies empirical filters based on our previous benchmarking efforts and expectation from the literature (see Methods). In brief, the filter retains SVs >= 250 bp, annotated with a single TE copy (single Repeatmasker hit ID) from members of the AluY lineage (Alu, N = 4,908), L1HS (LINE1, N = 704), SVA D, E and F subfamilies (SVA, N = 112) and HERVK from the HML2 lineage (HERVK, N = 31). HERVK candidates were further processed with a dedicated module categorizing the integrants into solo-LTR and proviral alleles (see Methods). Following the discovery phase, candidates were graph-genotyped with giraffe (Sirén Jouni et al. 2021) and only variants with long-read support were kept for final analysis (Figure S2). Additional polymorphisms within fixed (non-MEI) elements were reported: 10 within HERVK (7 solo LTR against provirus, and 3 with proviral copy-number variation) and 87 SVA VNTR-only events (tandem repeat expansions/contractions within fixed SVA loci) (Figure S2). Furthermore, 6 HERVK loci were initially split across 2 (N = 4) or 3 (N = 2) SVs each, and automatically reconciled by the GraffiTE HERVK classifier (Figure S2).

Using 1Mb bins, we observed MEIs loci distributed throughout the CHM13 reference genome with expected global depletions in some of the larger centromeric and pericentromeric regions although noticeable enrichments can be observed locally (Figure 2B). MEIs appeared also abundant in the proximal side of the extended MHC region (xMHC). The xMHC is known for its increased genomic variability (Trowsdale and Knight 2013), including CNV which is a putative source of false positives for MEI calling.

**Fig 2.**
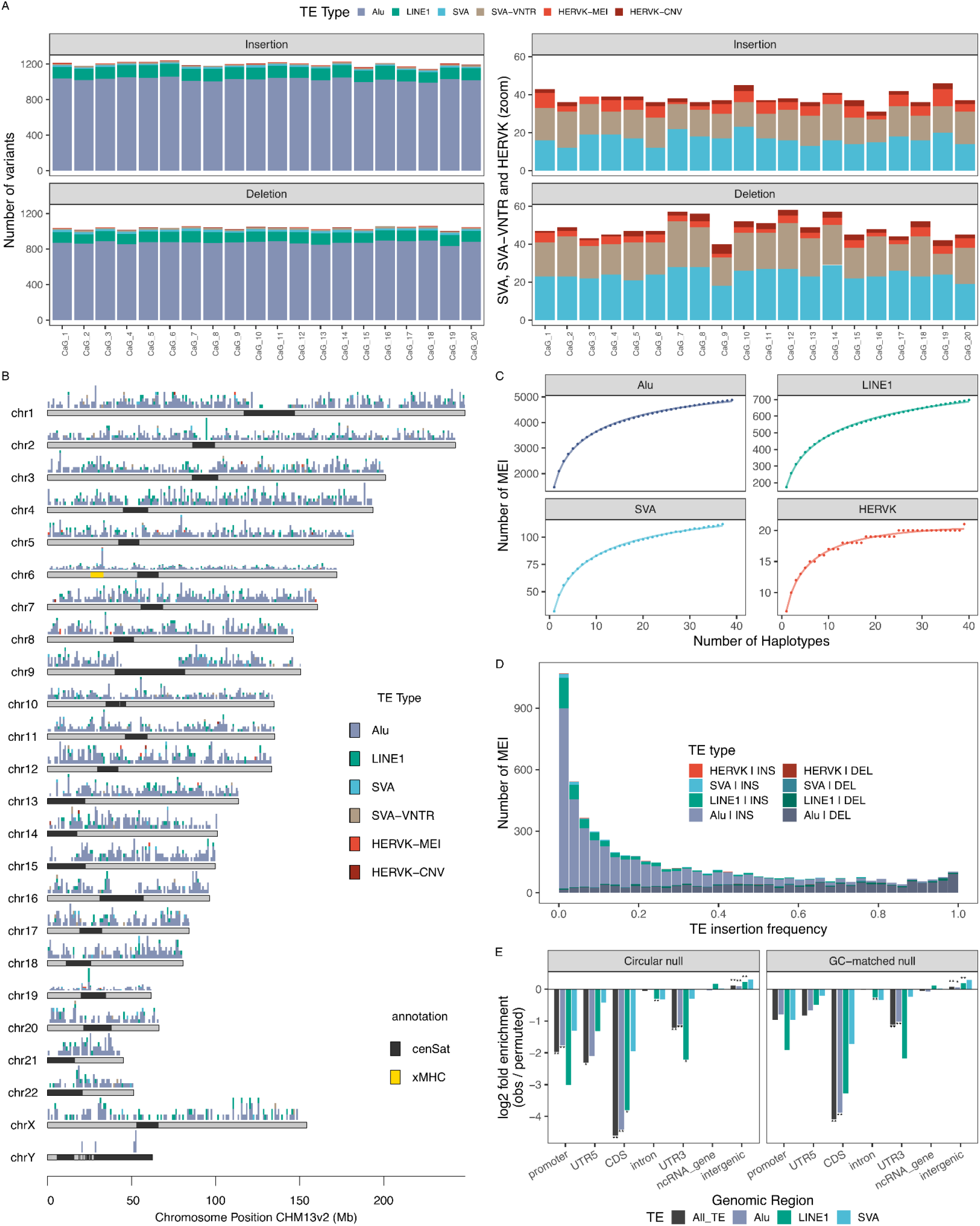
Annotation and population genetics of transposable elements in CaG. A) Distribution of TE-related variants among insertions (INS) and deletion (DEL) including MEI (Alu, LINE1, SVA), SVA-VNTRs (Variable Number of Tandem Repeat within reference SVA elements) and HERVK-CNV (Copy Number Variation of the proviral HERVK sequence) polymorphisms. B) Karyoplot representing the distribution of TE type among TE-related variants in bins of 1Mb. C) Discovery curves for each MEI type reported by GraffiTE over 40 haplotypes (graph genotypes). D. Insertion frequency spectrum stratified by MEI and SV type. E. MEI enrichments/depletions (log2 fold enrichment observed/permuted) across genomic regions, against a circular null (left; breakpoints of each chromosome shifted by one random distance) or a GC-matched null (right; random positions with the same flanking GC content). HERVK MEIs were excluded due to low sample size (N = 21). *: *P*-adj (B.H.) < 0.05, **: *P*-adj < 0.01.

To assess how much of the global TE variation is captured in the CARTaGENE cohort, we computed saturation curves for each of the 4 MEI types. Consistent with a small pangenome size, Alu, LINE1 and SVA MEIs keep being steadily discovered with additional samples; meanwhile MEI HERVK count distinctly showed early signs of saturation as new haplotypes are added (Figure 2C). A comparison of the GraffiTE MEIs with a joint SV call performed with minigraph-cactus (Hickey et al. 2023) on the CARTaGENE and HPRCv2 (Lucas et al. 2026) cohorts identified 314 MEIs (254 Alu, 52 LINE1, 7 SVA and 1 HERVK; 5.5% of GraffiTE MEI calls) private to the genome of these participants of French-Canadian descent (higher bound, see Methods), >95% of them singletons (299/314).

Next, we assigned the derived allele (polarized) of each MEI variant to compute the insertion frequency spectrum (IFS, Figure 2D). Due to their copy-and-paste transposition mechanisms, human MEIs are virtually homoplasy-free (Rishishwar et al. 2015; Xing et al. 2007), and thus the IFS is equivalent to an unfolded site frequency spectrum (SFS), where the insertion state (TE presence) represents the derived allele. Most segregating TEs, primarily insertions relative to CHM13, were found at low frequencies across the 20 individuals. Conversely, a smaller number of frequent variants, mainly drawn from deletions, showed an expected uptick at the end distribution. This u-shaped distribution of the IFS is consistent with the SFS computed on neutral regions of the human genome (N. Marchi and Excoffier 2020) supporting the paradigm that most MEIs are neutral (Cordaux et al. 2006; Arkhipova 2018). Furthermore, the IFS analysis confirms that including non-reference MEIs (deletions), often non-reported by alternative methods, is required in order to provide a complete picture of human population genetics parameters based on TE polymorphisms.

We further partitioned the genome based on MANE transcript annotation (Morales et al. 2022) to perform MEI enrichment analysis relative to gene positions (excluding HERVK due to low sample size, see Methods). We compared the observed counts with two permutation nulls: a circular null, in which all breakpoints of a chromosome are shifted by the same random distance (keeping their number and spacing), and a GC-matched null, in which each breakpoint is replaced by a random position with the same GC content in its flanking sequence. We added the second null because the L1 endonuclease, also used by Alu and SVA, cuts DNA preferentially at AT-rich motifs (Jurka 1997), and promoters and exons are GC-rich. We found MEIs globally depleted in promoter regions (defined as 1kb upstream of MANE transcription start site (TSS)), CDS and 3’UTR, principally driven by Alus; against a GC-matched null, only the CDS and 3’UTR depletions remained significant (Figure 2E). Pooled MEIs and Alus were also enriched in intergenic regions (9% and 7% above the circular null, 6% and 4% above the GC-matched null, respectively). LINE1 was depleted in CDS and 3’UTR (circular null only) but not significantly in promoters, and was reduced in introns and increased in intergenic regions, contrasting with SVAs that did not show deviation from the null expectation, which we hypothesize stems from their lower sample size (N = 106) (Figure 2E). MEIs were also depleted in the proximal and distal enhancer-like candidate cis-regulatory elements (cCREs) of ENCODE, including against a null matched for the GC content around each breakpoint (e.g., 345 breakpoints in distal enhancer-like elements against 641 expected), and in cCREs both active and inactive in blood (Fig S3; see Methods). Two processes could produce these depletions: (i) the preference of the L1 endonuclease for AT-rich target sites, which are less frequent in GC-rich promoters, exons and cCREs, and (ii) purifying selection against insertions that disrupt functional sequences. Experiments show that induced L1 insertions occur in all chromatin states, and are mainly constrained by local sequence preferences, whereas the distribution of older L1 copies was reshaped by selection (Flasch et al. 2019; Sultana et al. 2019). Taken together, these results support the well established role of purifying selection in shaping MEI distribution, with a likely smaller contribution of target-site preference.

### Characterization of complex alleles and insertion structure of TE variants

To resolve HERVK loci whose alleles differ in the structure of the element rather than in its presence, we implemented a HERVK classifier in GraffiTE (see Methods). A full-length HERVK insertion (provirus) consists of an internal coding region flanked by two long terminal repeats (LTRs) that are identical at the time of insertion. When the two LTRs recombine, the element loses its internal region and one LTR and becomes a solo LTR (Thomas et al. 2018). The solo LTR and the provirus can then segregate at the same locus; we call such a locus dimorphic when every haplotype carries one of the two forms. Conversely, unequal exchange between the LTRs of two chromatids can occasionally produce two or more proviruses in tandem (proviral copy number variation) (Hughes and Coffin 2004; Pasternack et al. 2025). A single locus can segregate several of these states, and in an SV call set each of them appears as a separate insertion or deletion of HERVK sequence. Without the classifier, we would count the difference between a solo LTR and a provirus as an MEI, although no transposition separates the two. The classifier instead counts each allele in proviral units (empty site, solo LTR, one provirus or several in tandem) and merges the records of a locus into one. We identified 31 HERVK (HML-2) loci across the 20 samples (Figure 3A). Twenty-one are MEIs, at which a null allele segregates: 16 segregate a solo LTR against the empty site, 4 a provirus against the empty site, and one, K134 at 12q13.2, all three states. The other ten carry the element on every haplotype we called (Figure 3A). Seven are dimorphic, segregating a solo LTR against a provirus. Three vary in proviral copy number: K108 at 7p22.1 (one, two and three units), K109 at 6q14.1 (a solo LTR, one and two units) and K42 at 12q24.33 (one and two units). Fifteen of the 31 loci carry a published HML-2 name: ten are in the HML-2 catalogue published by Subramanian et al. 2011, among them K106, K107, K108, K109, K113 and K115, and five were reported in the Neanderthal or Denisovan genomes (Agoni et al. 2012; E. Marchi et al. 2014) . We identified K115 by sequence alignment, as liftOver was unable to map its hg19 coordinate to CHM13. Of the 31 loci, 24 are reported by Wildschutte et al. 2016, and at the 12 loci where they report an element form, it matches ours: 10 solo LTRs and 2 proviruses (Table S1). Of the seven loci they do not report, four segregate a null allele in our cohort. Conversely, 18 loci they genotyped are absent from our call set. Fourteen of these have an allele frequency below 0.025 in their data. Insertion frequencies at the 17 MEI loci present in both studies correlate with theirs (Pearson r = 0.74; Spearman ρ = 0.82, P = 5 × 10⁻⁵; Figure 3B).

**Fig 3.**
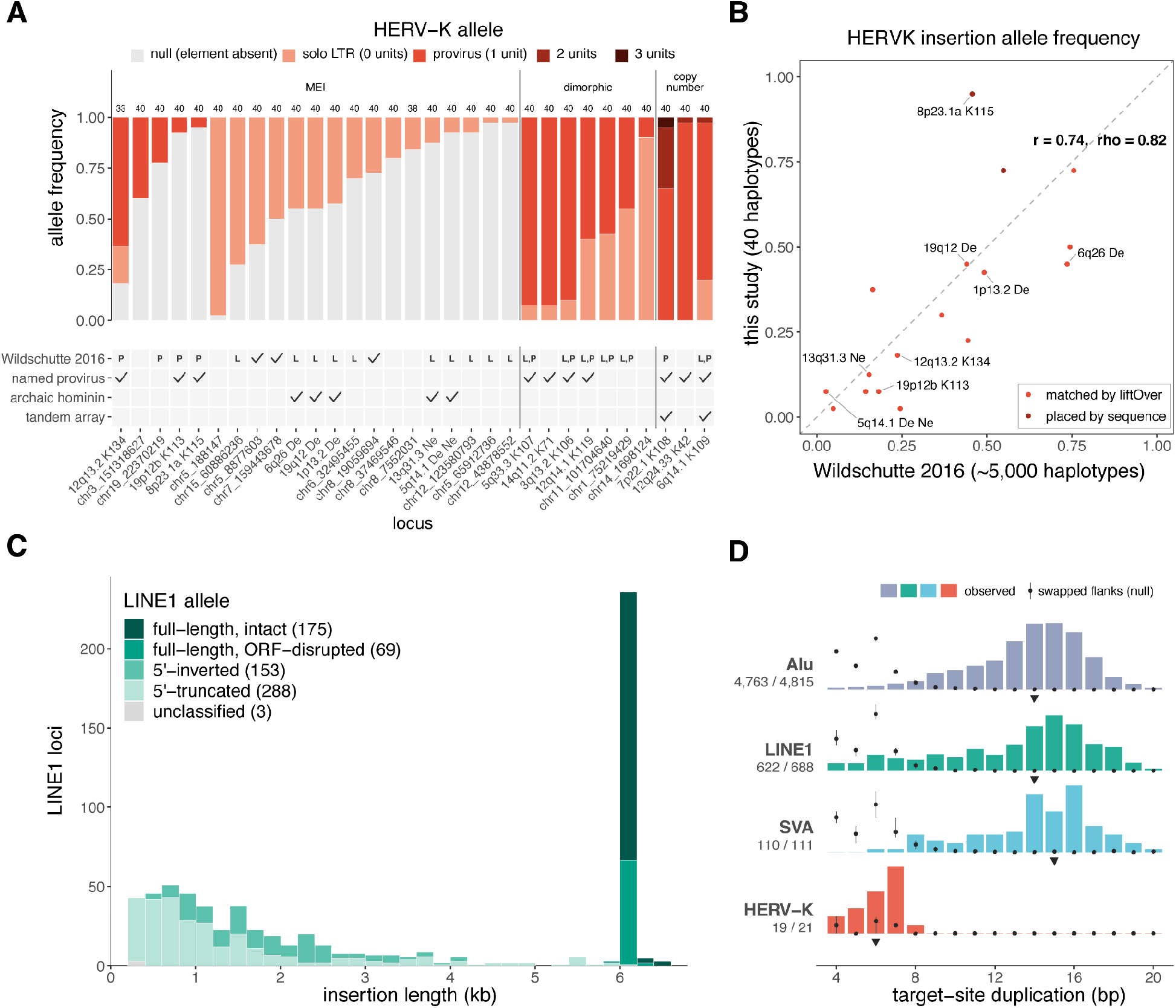
Allele and insertion structure resolved by GraffiTE. A) HERVK (HML-2) allele frequencies at 31 loci over 40 haplotypes, grouped as MEI (a null allele segregates; 21 loci), dimorphic (solo LTR against provirus; 7) and proviral copy number (alleles differ in the number of proviral units; 3). The number above each bar gives the haplotypes called. Copy-number loci use genotypes from the haplotype-resolved assemblies, because GraffiTE withholds their graph genotypes; all others use graph genotypes. Below, a mark means the studies of that row report the locus. Rows: Wildschutte et al. (2016), a survey of polymorphic HML-2 insertions in about 2,500 individuals; named provirus, a locus named in the HML-2 catalogue of Subramanian et al. (2011); archaic hominin, an insertion that Agoni et al. (2012) or Marchi et al. (2014) report in Neanderthal or Denisovan genomes; tandem array, a locus where Hughes and Coffin (2004) or Pasternack et al. (2025) report tandem proviral copies. On the Wildschutte row, P is a provirus and L a solo LTR in their data, L,P both forms, and a tick a locus they report without element form. Loci with a published HML-2 name carry it; others are labelled by CHM13 position. B) Insertion frequency at the 17 MEI loci present in both studies, this study (40 haplotypes) against Wildschutte et al. (about 5,000 haplotypes), with y = x. C) Length of the 704 LINE1 MEIs, filled by structural state. Full-length means the element aligns to L1.3 from its first 50 bp over at least 95% of its length, and intact means ORF1 and ORF2 are both open. 5′-inverted elements carry the inverted 5′ segment left by twin priming; 5′-truncated elements are 5′-incomplete without one. Internally deleted elements start within the first 50 bp of L1.3 but cover less than 95% of it. D) Target-site duplication (TSD) length by family. Bars: share of each family’s insertions with a TSD of each length. Triangles: the median. Labels: insertions with a TSD over those tested. Grey points: the same search on windows swapped between insertions of the same family, strand and SV type, where no duplication spans the junction (mean of 10 replicates, whiskers lowest to highest). Bars and points share one scale within a family. HERVK is shown at its MEI loci only.

Leveraging the detailed annotation from GraffiTE, we characterized each LINE1 according to sequence completeness and organisation (Figure 3C). We found that 64.2% of the LINE1 MEIs bear a characteristic 5’ truncation caused by the premature termination of the target primed reverse transcription (TPRT) process (Ostertag and Kazazian 2001; Meyer et al. 2010). Of them, 155 (22.0% of LINE1 MEIs) were also reported to carry a 5’ inversion, which occurs through twin priming and derived mechanisms (Ostertag and Kazazian 2001; Meyer et al. 2010), consistent with our previous estimate in 94 haplotypes of the HPRCv1 cohort (18.7%, 451/2413) (Groza et al. 2024). While 248 LINE1 were full-length, only 175 appeared intact, and thus potentially active, upon inspection of their ORF.

GraffiTE reports the target-site duplication (TSD) of each insertion. We found a TSD at 4,856 of 4,908 Alu (98.9%), 638 of 704 LINE1 (90.6%) and 111 of 112 SVA (99.1%) MEIs, with median lengths of 14, 14 and 15 bp (Figure 3D), closely matching the 15 bp expectation for L1-mediated insertions (Szak et al. 2002; Kojima 2010). HERVK MEIs carried shorter TSDs (19 of 21; median 6 bp), consistent with the 5 to 6 bp reported for HML-2 (Kahyo et al. 2017). To measure how often the search finds a TSD by chance, we ran it on windows swapped between insertions of the same family, strand and SV type, where no duplication spans the junction (Methods). The search found a TSD of 9 bp or more in 1.7% of swapped Alu, LINE1 and SVA pairs, against 91.4% of real insertions. Among LINE1, 5′-truncated insertions lacked a TSD more often than full-length ones (18% against 1.6%) and, when found, it was more often 8 bp or shorter (30% against 6.6%), as reported for 5′-truncated L1 (Kojima 2010).

#### GraffiTE surveys the TE panepigenome

Next, we turned to quantifying the methylome in newly characterized MEIs (Methods). To do so, we created a genome graph that represents MEIs across the 20 diploid genomes, enabling us to count the number of CpGs involved in each TE family (Methods). In total, we counted that the MEI panepigenome consists of 144,748 CpGs. Of these, 104,782 are in Alus, 27,443 are in LINE1, 11,808 in SVAs and 715 in HERVKs (Fig 4A). To determine whether the MEI panepigenome is closed and will eventually plateau in growth, we applied Heaps law with quadratic correction (Tettelin et al. 2005; Fontanelli and Li 2025; Heaps 1978). For Alu, LINE1, and SVA, we found that the panepigenome is open but slowing relative to Heaps’ Law (Table S2, Fig S4, Methods). The HERVK component of the panepigenome grows the slowest, which is consistent with the relative activity of families in the human genome. In a sufficiently large pangenome, we expect the MEI panepigenome growth rate to be determined by the rate of transposition of each family.

**Fig 4.**
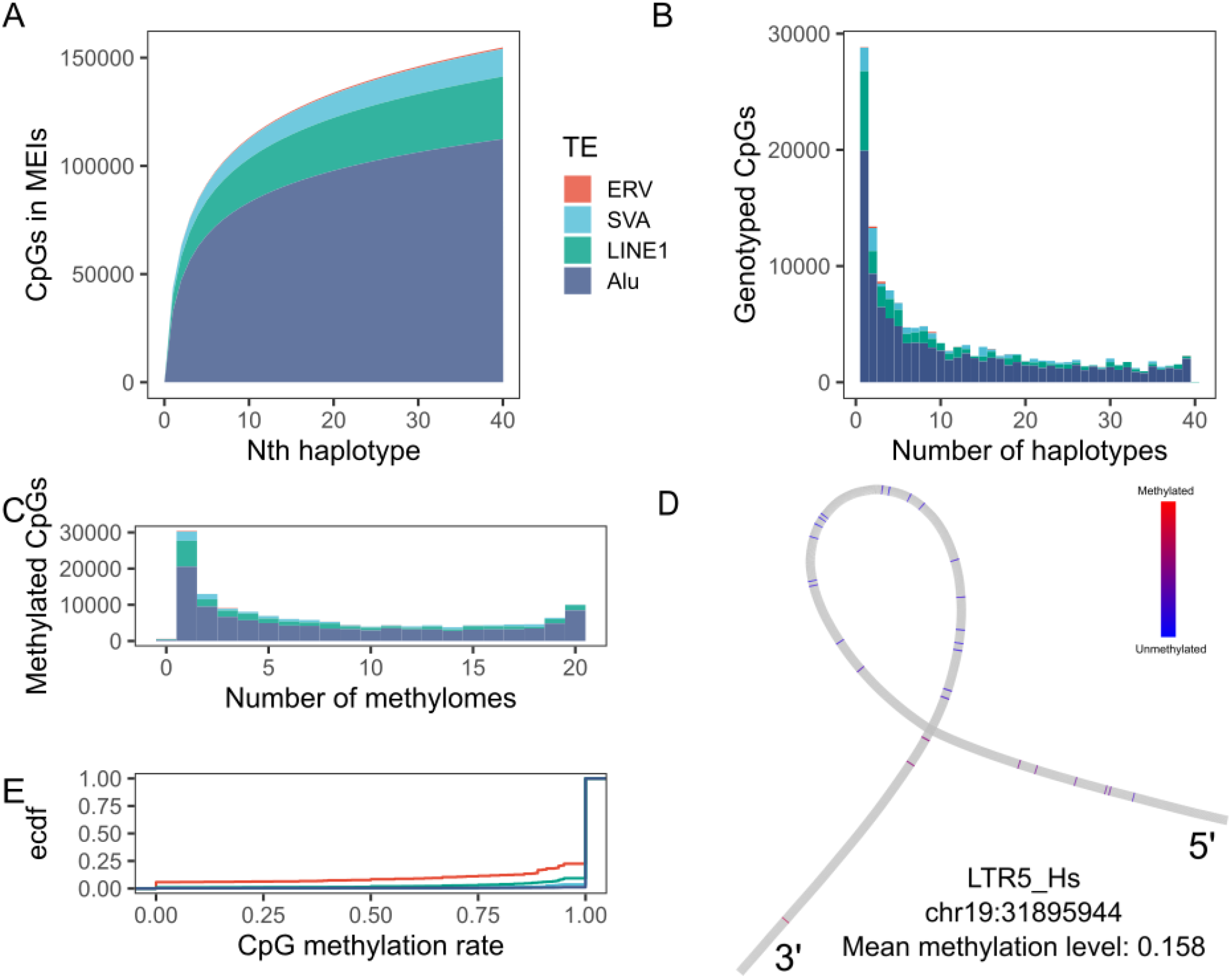
A) Number of CpGs involved in MEIs for each additional explored genome, stratified by family. B) CpG allele counts stratified by TE family in the 40 CARTaGENE haplotypes. C) Methylated CpGs count in the 20 CARTaGENE methylomes stratified by the TE family. D) Example HERV LTR5_Hs containing 19 hypomethylated CpGs. The CpGs in the immediate vicinity are relatively more methylated. E) Empirical cumulative distribution of the methylation rates of CpGs across TE families.

When looking at the allele count of each CpG, 28,830 were singletons, while 6,617 had an allele frequency of at least 90% (Fig 4B). Next, we wanted to find epigenetic variation of CpGs within MEIs. Thus, we mapped the 20 matching methylomes to the TE variation. Across all CpGs, only 602 were never methylated. Meanwhile 21,148 CpGs were methylated in at least 18 (90%) of methylomes (Fig 4C). However, the CpG methylation count is limited by its allele count. When correcting for allele count to obtain methylation frequency (MF), we found 142,141 CpGs with MF ⩾ 90% and only 631 CpGs with MF ≤ 10% (Fig S5). Thus, we conclude that CpGs in MEI are overwhelmingly methylated, providing a fully independent validation of the observation reported by (Zhuo et al. 2026) in the draft human pangenome cohort that is in line with prior expectations (Yoder et al. 1997).

Individual CpGs and their associated methylation values can be visualized with Bandage and GraffiTE. We show an LTR5_Hs HERVK insertion (allele count: 13) with an average methylation level of 15.8% over its 966 bp length. The CpGs within the MEI exist in a relatively hypermethylated immediate genomic context (chr19:31895944, Fig 4D). The epigenetic distinctness of such events compared to their vicinity could highlight TEs with potential regulatory properties.

Then, we asked if the methylation level of individual CpGs is the same across MEI families (Fig 4E). Here, we found that the distribution of methylation levels of CpGs in LINE1s (mean: 72.2%) and HERVKs (mean: 73.3%) is skewed towards lower values relative to Alus (mean: 87.1%) and SVAs (mean: 94.5%).

### TE methylation decreases with age and genomic context

GraffiTE estimates methylation levels in fixed TEs at the same time. These estimates are in line with established methods such as pb-cpg-tools (R^2^: 0.9793, Fig S6, Methods). Using this feature, we compared the methylation level of the MEIs above to the methylation level of fixed TEs (Fig 5A). We found that polymorphic Alu, LINE1 and SVA TEs feature higher methylation levels than their fixed counterparts (overall effect: 12.7%, P <0.0001, Fig 5A, Fig S7, Tables S2-S3), confirming a result previously observed in Groza et al. 2025. The presence of hypomethylated MEIs is intriguing since they could be copies that escaped repression. Indeed, we identify 480 Alus, 67 LINE1s, 12 SVAs, and 2 HERVK in hypomethylated states (Methods).

**Fig 5.**
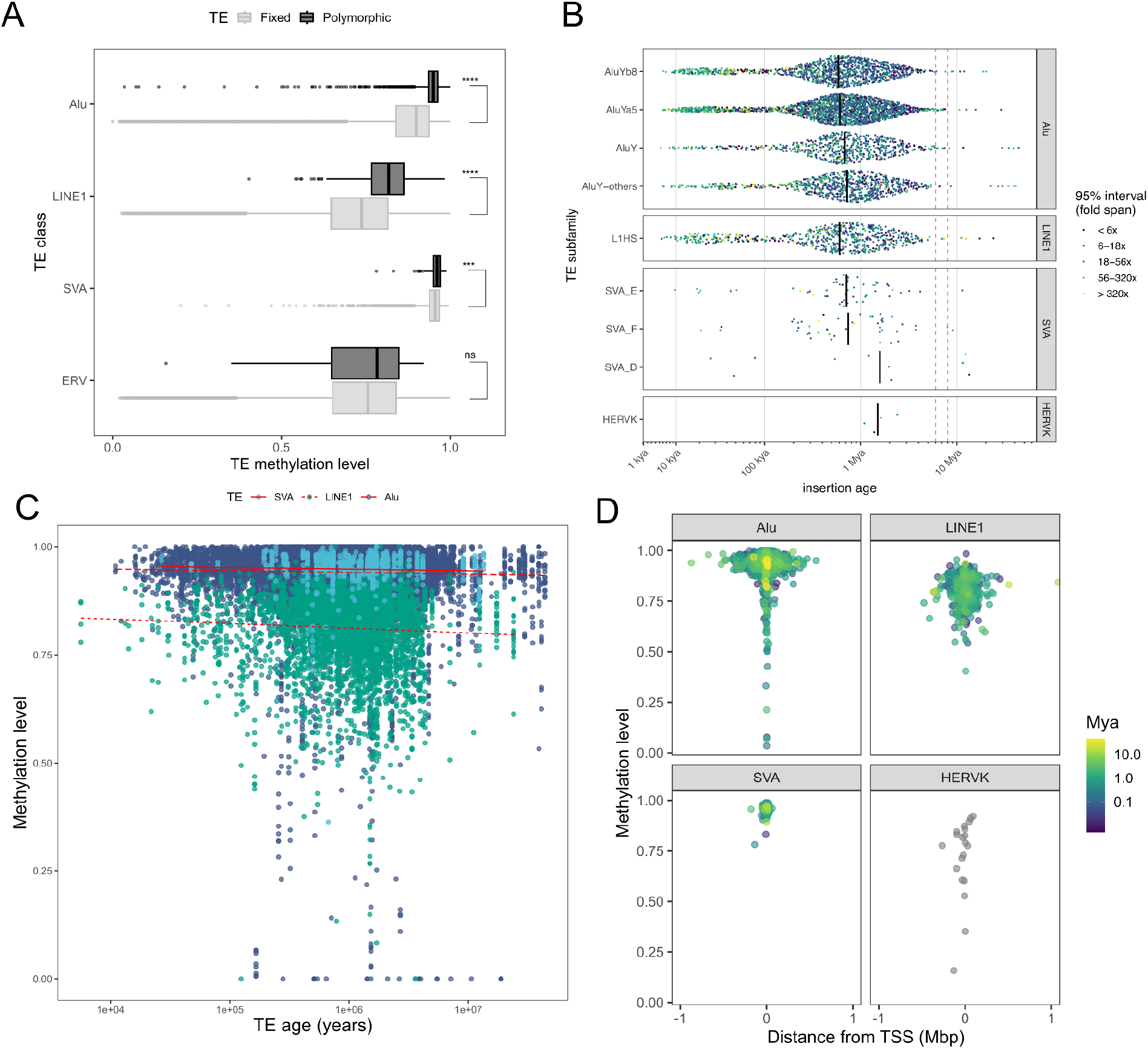
A) Average CpG levels methylation across fixed and MEI annotated by GraffiTE. B) Age dating of each MEI, stratified by TE class and subfamily. The red lines denote an age range dating the split from Chimpanzee. HERVKs are subset to those that respect the assumptions of the dating method. C) Beta regression estimating the effect of age on the methylation levels of each MEI. D) Distance of each MEI relative to transcription start sites, color scaled by their age and stratified by TE class.

We hypothesized that older MEIs may be demethylated relative to younger MEIs. To estimate the effect of age, we first dated polymorphic MEIs based on an ancestral recombination graph (ARG, Table S5) (Methods). We inferred that Alus are 1.04 Mya old on average, LINE1 1.06 Mya and SVAs 1.51 Mya (Fig 5B). These ages are of the same order as earlier estimates based on sequence divergence, which date polymorphic AluYa5/8 elements to 1.63 Mya on average (Arcot et al. 1996) and the AluYb8 and AluYb9 subfamilies to 2.39 and 2.32 Mya (Carter et al. 2004). Likewise, the mean LINE1 and SVA ages predate the emergence of the subfamilies active in humans: the L1 Ta family arose approximately 4 Mya (Boissinot et al. 2000), and the SVA_E and SVA_F subfamilies are restricted to the human lineage (Wang et al. 2005).

To test the effect of MEI age on DNA methylation levels in blood, we fitted a beta regression model jointly over all MEI and included family, donor age, GC content, the local recombination rate, genomic context and blood cell type composition as covariates (Fig 5C, Table S6, Methods). According to the model, the expected methylation levels of MEIs are determined, among other factors, by their family (P_LINE1_ < 2.23x10^-16^, P_SVA_ < 6.44x10^-7^,) and genomic context (P_promoter_ < 0.0048). As expected, donor age also had a statistically significant methylating effect (0.0022 log-odds per donor year, P < 5.58x10^-4^) on MEIs. When correcting for blood cell type composition, a higher basophil proportion had a global demethylating effect (P < 3.58x10^-4^). Interestingly, increasing MEI age retains a statistically significant demethylating effect (-0.069 log-odds per log10 years, P < 6.08x10^-19^). More precisely, we expect an average LINE1 insertion to demethylate from 86.4% to 80.8% over the first Mya. Meanwhile, the methylation of non-autonomous MEIs remains stable over time, with Alus decreasing from 95.6% to 93.5% and SVAs decreasing from 96.2% to 94.3% over the same time interval. We also tested alternative sets of dates (Albers and McVean 2020) obtained based on a molecular clock (MC, Table S7) and recombination clock (GEVA, Table S8, Methods). The MC dates recapitulate the ARG association (-0.059 log-odds per log10 year, P < 8.14x10^-16^) while GEVA finds a similar albeit weaker association (-0.034 log-odds per log10 year, P < 8.90x10^-8^) . We expect allele count to be a noisy estimator of age (Figure S8) and should recapitulate a portion of the effect. Indeed, MEIs demethylate as they near fixation (-0.072 log-odds, P < 8.32x10^-10^, Table S9, Methods). Given that LINE1s are autonomous and demethylate the most with age, we categorized LINE1 according to their predicted activity potential (Methods) and tested its impact on methylation together with age. In this model, LINE1 age is no longer statistically significant while LINE1 activity potential is associated with increased methylation (log-odds 0.35, P < 2.23x10^-16^, Table S10). This supports the hypothesis that methylation is sustained in LINE1s with transposition potential, while LINE1s that lose that ability with time are free to demethylate.

We also confirmed the interplay between methylation, age and the location of MEIs in the genome that was suggested by our modelling. When stratifying MEIs between promoter, exon, intronic, intergenic and non-coding RNA regions, we find that the average methylation is lowest in promoters (mean: 72.5%), while other regions range from 91.6% to 93.4% (Fig S9). Indeed, demethylated MEIs tend to concentrate near transcription start sites (Fig 5D). MEIs in promoters are older (mean: 1.11 Mya, n = 11) than those in introns (mean: 819 kya, n = 2018, Fig S10) but younger than those in intergenic regions (mean: 1.19 Mya, n = 2839). MEIs in exons are very rare (n = 2). However, the demethylated MEIs in promoters are not necessarily the oldest (Fig 5D). Thus, any relationship between age and methylation is contingent on the local regulation of the epigenome. For instance, an ancient MEI in a promoter may be highly methylated because the downstream gene is not expressed in blood. Meanwhile, a young TE is demethylated because it is inserted in the promoter of an active gene. In conclusion, age is associated with slow demethylation, but extreme demethylation of MEIs is also contingent on genomic and regulatory context in humans.

## Discussion

We demonstrated that GraffiTE v1.1 leverages the full potential of LRS to genotype SVs, annotate MEIs and map the epigenetic state of TEs in the genomes with matching methylomes of 20 Quebec residents of French-Canadian descent. Due to its transparent design, we could combine GraffiTE v1.1 with assembly validation tools like HMM-Flagger to remove false positives, improve the quality of genotypes and obtain more robust allele frequencies.

Among MEI genotyping pipelines, GraffiTE provides the richest set of features by annotating LINE1s with 5’ truncation and 5’ inversion, by classifying HERVK alleles as empty site, solo LTR, provirus or proviral copy number, and by flagging VNTR polymorphisms within SVAs. Furthermore, we provide an improved TSD module whose sensitivity is tested against a null distribution and precision is validated against curated human datasets. Although this study focuses on human polymorphisms and highlights the performances of its novel dedicated filter, GraffiTE is applicable to any eukaryote system.

Unlike other similar tools, our approach does not discard the epigenetic data provided by LRS, or does not restrict its readout to non-reference MEI (insertions). Instead, we map this information to the entire genome, including MEIs. In this effort, we already identified hypomethylated MEIs and visualized their epigenetic state at basepair resolution within variation graphs, a unique distinction that critically enhances mappability at and around structural variants (Groza et al. 2020). Therefore, GraffiTE identifies outlying MEIs of potential biological interest. Critically, these results allowed us to detect a demethylating effect of insertion age upon mobile elements, and also related to LINE1 activity potential. We note the challenges associated with dating MEIs based on only 40 haplotypes. However, the ranking of the variants through time should still be sufficiently reliable to detect the effect of age on methylation, if one exists. The exact magnitude of the demethylating effect will be best confirmed and explored in the third planned release of the HPRC genomes and methylomes. Furthermore, our epigenetic annotation workflow enables deeper annotation that could point to MEIs with regulatory activity in specific biological contexts. In fact, we already observed several demethylated MEIs near transcription start sites.

In the past, it was difficult to map transcription factor and histone epigenomes to MEIs due to the challenging mappability of short-read assays like ChIP-seq to TEs. Today, emerging techniques based on stencils can annotate highly mappable long-reads with epigenomic features such as chromatin accessibility, bound transcription factors and histone modifications. GraffiTE v1.1 is positioned for these possibilities by remapping any available base modification to MEIs. In Fiber-seq, GraffiTE could map the methylated adenine to MEIs and reveal their chromatin accessibility status. In this manner, GraffiTE v1.1 pioneers future epigenomic studies in MEIs that go beyond CpG methylation.

## Methods

### Sample selection in CARTaGENE

To select CaG participants for long read sequencing, we started with a pruned set of variants with minor allele frequencies 5% from the ≥ 29,337 CaG participants with SNP array data available. We made sure no participants were related to at least the 4th degree. Based on the CaG clusters determined by UMAP and HDBSCAN from (McClelland et al. 2026), we excluded participants who were not in clusters 13 or 14, corresponding respectively to the Saguenay and the French-Canadian cluster, participants for whom their first learned language was not French, and participants for whom at least one of their four grandparents was not born in Canada. We then applied the K-means algorithm to cluster the participants in 10 clusters using the first 10 principal components calculated from our pruned set of variants. We then randomly selected 2 participants per cluster for whom short-read whole-genome sequencing data were available.

#### Sequencing of CARTaGENE genomes

PacBio libraries were prepared according to manufacturer specifications, following protocol "Preparing whole genome and metagenome libraries using SMRTbell® prep kit 3.0" (PN#102-166-600 REV05 DEC2024). DNA Samples were sheared using the Diagenode Megaruptor® 3 and the Megaruptor 3 Shearing Kit (cat#E07010003), with an input of 5 ug in a final volume of 130 uL (final concentration 38.46 ng/µL), followed by a AMPure PB (cat# 100-265-900) bead cleanup (1.0X v/v) to concentrate the samples in 46 µL. Then, using Pacific Biosciences SMRTbell® prep kit 3.0 (cat# 102-141-700), the sheared DNA was end repaired, A tailed and ligated to the SMRTbell barcoded adaptors (SMRTbell adapter index plate 96A cat# 102-009-200). Following this, the library was exonuclease treated and underwent an AMPure PB 35% volume/volume size selection with a ratio bead to template of 3.1X, yielding between 0.712 to 1.579 ug of final library products. The prepared library were then annealed to the PacBio sequencing primer bound to the Revio polymerase (cat#102-817-600) and cleaned with the PacBio SMRTbell cleanup beads (cat# 102-158-300). The bound complex was then loaded in the sequencer at a loading concentration of 300pM where the library was sequenced using the Revio sequencing plate (cat# 102-587-400) on one Revio SMRT Cell (cat# 102-202-300) per library using a PacBio Revio sequencer with a 30h movie time. Top up SMRTcells (to reach 30X genome coverage) were sequenced using Revio SPRQ polymerase (cat# 103-496-900) and Revio SPRQ sequencing plate (cat# 103-512-700). Primary analysis was done on instrument using SMRT Link software version 13.3.0.253824.

#### Base and methylation calling

For assembly-based methylation estimates, we employed pb-cpg-tools’ aligned_bam_to_cpg_scores with --pileup-mode count and where --modsites-mode is the sample’s *de novo* assembled hap1 and hap2.

#### Assembly and validation of CARTaGENE genomes

To assemble the genomes, we used hifiasm (Cheng et al. 2021), as implemented in the cgroza/hifiasm-nf pipeline on GitHub: `nextflow run cgroza/hifiasm-nf.nf --design design.csv.` The result was 40 *de novo* pseudo-haplotype assemblies.

To validate the correctness of the resulting genome assemblies, we remapped the PacBio data to the assemblies and annotated erroneous, duplicated and collapsed sequences with HMM-Flagger , as implemented in the cgroza/HMM-Flagger pipeline on GitHub using the hifi presets: nextflow run nf-HMM-flagger/main.nf --samples flagger.csv.

To evaluate the span of CHM13 covered by our assemblies, we mapped both haplotypes to CHM13 with minimap2 (Li 2018) and obtained the percent coverage with `samtools coverage` (Li et al. 2009).

#### Structural variation calling

We called structural variation (SVs) with PAV (Ebert et al. 2021) to obtain insertions and deletions relative to CHM13 (Nurk Sergey et al. 2022), together with the supporting genome intervals in the CARTaGENE assemblies (`nextflow run cgroza/GraffiTE --pav`). We removed any SVs supported by unreliable assembly regions as annotated by HMM-Flagger. We merged the SVs into a single multi-sample VCF with truvari (English et al. 2022).

#### Transposable element annotation with GraffiTE

To annotate SVs, we used an updated version of the GraffiTE workflow along with the consensus sequence set of human TE families available in Dfam 3.9 (see below). Note that the recent release 4.0 of Dfam does not affect the human library. This novel version of GraffiTE (v1.1.0.0) pipeline merges SVs across samples with truvari (English et al. 2022), extracts variant sequences, annotates these with RepeatMasker (Hubley et al. 2016) and outputs an annotated VCF file and a corresponding graph genome. We used the following GraffiTE command: nextflow run cgroza/GraffiTE --TE_library human_DFAM3.9.fasta --vcfs vcfs.tsv --reference CHM13.fa --human. The --human flag, new in GraffiTE v1.1, keeps SVs of at least 250 bp annotated as AluY, LINE1 (L1HS), SVA (SVA_D, SVA_E and SVA_F) or HERVK of the HML-2 lineage, and classifies the HERVK alleles (see below).

### GraffiTE 1.1

We generated the call set with the development code of GraffiTE 1.1 while we finished the public release (v1.1.0.0). When new experimental features were added to the production pipeline, we curated the existing records to match the behavior of the release version. In the pipeline, GraffiTE annotated each SV with RepeatMasker 4.2.4 and ULTRA 1.0.0 and kept the 132,511 SVs whose sequence was more than 80% covered by repeats. We genotyped the samples with vg giraffe and vg call (vg 1.77.0). The --human filter and the HERVK classifier selected the records we analyse (next sections). Outside the pipeline, we corrected the sex-chromosome genotypes, removed loci without read support, took discovery genotypes at three HERVK loci, added 114 records that only the release keeps (next section), and searched for target-site duplications (TSDs) with a search we redesigned after the runs and then added to the release. The headers of the VCFs we provide name each run and each step done outside the pipeline. We used default parameters unless stated.

### GraffiTE v1.1 --human filter

The novel --human mode aims at keeping the records that can represent a polymorphic copy of a mobile element family active in humans. A record passes when six conditions are met. (1) Its RepeatMasker class and subfamily, matched by name prefix, are SINE/Alu with AluY, LINE/L1 with L1HS, Retroposon/SVA with SVA_D, SVA_E or SVA_F, or LTR/ERVK with HERVK-int, HERVK, LTR5_Hs, LTR5A or LTR5B; SVA records annotated only by their VNTR (class Simple_repeat) pass under the SVA list (these regular expressions can be expanded or restricted by the user). (2) |SVLEN| is at least 250 bp. (3) ULTRA tandem repeats cover less than 60% of the variant sequence; VNTR-only SVA records are exempt. (4) The record has a single RepeatMasker hit, or up to three when an HERVK-int hit lies beside an SVA hit and the variant is at most 10,500 bp. The SINE-R domain of SVA derives from the HERVK LTR, so RepeatMasker can assign part of a provirus to SVA, and it can split the internal region of a provirus in two. (5) A single-hit Alu, LINE1 or SVA record ends in a poly(A) tail: a run of at least 8 bp, at least 80% A, within 5 bp of the 3′ end of the variant sequence, with or without the TSD copy at that end, or the same run of T at its 5′ end when the element lies on the minus strand. HERVK and VNTR-only SVA records are exempt. (6) FILTER is PASS. In this cohort the filter kept 5,933 of the 132,511 repeat-annotated SVs: 4,970 Alu, 718 L1HS, 115 SVA, 93 VNTR-only SVA and 37 HERVK records. Our runs looked for the poly(A) tail only in the variant sequence as called and did not keep 114 of these records (96 Alu, 17 L1HS and 1 SVA). We added them to the call set.

### Genotyping and sex chromosomes

In the pipeline, GraffiTE aligned the PacBio HiFi reads of each sample to its graph with vg giraffe (--parameter-preset hifi) and genotyped each sample with vg call. GraffiTE runs vg call with these defaults: -a, which reports a genotype at every snarl so that a sample lacking the variant gets an explicit 0/0 call; -A; minimum allele and site support of 2 and 4 reads (-m 2,4), below which calls are flagged lowad and lowdepth; and ploidy 1 on chrX and chrY for every sample (-R chrX:1,chrY:1). GraffiTE then split, merged and annotated the per-sample calls with bcftools v1.16 (norm -m-, merge -m none, annotate), giving 199,268 records.

With ploidy 1 on chrX, vg call genotyped females as haploid on chrX. Outside the pipeline, we replaced those genotypes. We assigned sex from the share of non-pseudoautosomal chrY records with a called genotype in an earlier, diploid genotyping run of the same samples against the same graph: 0.000 to 0.015 in 10 females and 0.661 to 0.730 in 10 males. Autosomal genotypes are identical between the two runs at all 112,098 called genotypes of the 5,637 loci they share, and alleles are identical at all 154 shared sex-chromosome records. We replaced every female chrX genotype with the call of the diploid run. Males stay haploid on chrX and chrY outside the pseudoautosomal regions (CHM13 chrX PAR1 1-2,394,410 and PAR2 153,925,834-154,259,566; chrY PAR1 1-2,458,320 and PAR2 62,122,809-62,460,029). We took the two chrX PAR1 loci from the diploid run in all 20 samples and removed the five chrY PAR1 records, because PAR sequence lies on both the chrX and chrY paths of the graph and its reads divide between them. No female was called at any of the 10 male-specific chrY loci. We set FORMAT/SXF = 1 on each replaced genotype.

### Curated call set

We kept the genotypes of the records that passed the --human filter. Of the 5,896 Alu, LINE1 and SVA records it kept (VNTR-only included), nine had no genotype (1 AluY, 2 L1HS, 6 VNTR-only SVA), and we removed the five chrY PAR1 records (above). We removed a locus only when no sample carried an ALT allele, which applied to 71 loci. To these 5,811 records we added the 31 HERVK locus records (see HERVK allele states), giving the 5,842 records of the analysis VCF. We set aside the 87 VNTR-only SVA records in analyses that treat a record as an insertion, leaving 5,755: 4,908 Alu, 704 LINE1, 112 SVA and 31 HERVK loci, 21 of which segregate a null allele.

### Target-site duplications

We searched for TSDs outside the pipeline, with the redesigned search now in the release, on the 5,933 records that passed the --human filter, except four added records with no ALT allele in any sample (5,929 records). For each record we built two 60 bp fragments. L holds the 30 bp of reference that end at the anchor base (the reference base before the variant, which the VCF record repeats in REF and ALT), then the first 30 bp of the variant sequence (ALT for an insertion, REF for a deletion). R holds the last 30 bp of the variant sequence, then the next 30 bp of reference. We listed every exact match of 4 to 20 bp shared by L and R. A TSD sits at the two junctions of the variant. Depending on where the caller placed the variant, its two copies both begin at a junction (one at the first variant base, the other just after the variant) or both end at one (one at the anchor base, the other at the last variant base). We scored each match by the mean distance, in bases, of its two copies from the nearer of these two arrangements; a score of 0 is a match exactly at the junctions. Short matches arise by chance more often, so a candidate shorter than 6 bp competed only at a score of 0.5 or less. The best candidate passed at a score of 5 or less. We report TSDs for the 5,745 MEI records, using for each HERVK locus its record with a null allele.

We estimated chance matches by drawing a cross-locus swap null distribution. Each record kept its own L fragment and took the R fragment of a donor with the same family (LINE1 split into full-length, 5′-inverted and 5′-truncated), RepeatMasker strand and SV type, never itself or a record within 10 kb on the same chromosome. We drew donors as a random permutation within each stratum and ran the unchanged search on 10 replicates; four LINE1 records outside these three classes had no donor. We report, per family and TSD length, the share of records with a passing call (mean and range over replicates).

### Structural classification of MEIs

#### LINE1 insertions

We classified the 704 LINE1 MEIs of the call set (521 insertions, 183 deletions) by structure with our Python package l1activity v0.1.0 (Python 3.13.7, pysam 0.23.3, Biopython 1.86); the package is distributed in the paper/CaG2026 directory of the GraffiTE repository. We took each element from the ALT (insertions) or REF (deletions) sequence without its anchor base and reverse-complemented elements on the minus strand. We aligned each element to L1.3 (GenBank L19088, 6,059 bp), a full-length L1HS that retrotransposes in cultured cells (Dombroski et al. 1993; Sassaman et al. 1997), with minimap2 v2.30 (-x asm20 --cs), and called it full-length when the alignment started within the first 50 bp of L1.3 and covered at least 95% of it. We derived the ORF1p (338 aa) and ORF2p (1,275 aa) reference proteins from L1.3, translated sense-strand open reading frames of at least 300 nt with EMBOSS getorf v6.6.0, and assigned them to either protein with blastp (BLAST+ v2.16.0, maximum E-value 1e-10). We scored an ORF as intact when one peptide covered at least 90% of the reference protein, ended within 3% of each of its termini, and shared at least 95% amino-acid identity with it. We took 5′ inversions from the GraffiTE VCF’s L1_5PINV field, which GraffiTE sets when an L1 RepeatMasker hit consists of a minus-strand fragment followed by a plus-strand fragment. We assigned each locus to the first class it matched: 5′-inverted (155 loci); full-length with both ORFs intact (175); ORF2 intact with ORF1 disrupted (25); full-length with ORF1 or ORF2 disrupted (57); 5′-truncated, meaning an alignment start beyond position 50 of L1.3 whatever the ORF status (288); and internally deleted, meaning an alignment that starts within the first 50 bp of L1.3 but covers less than 95% of it (1). An inverted element with an intact ORF2 would count as ORF2-intact; none had one. Three elements of 252 to 292 bp did not align to L1.3 and remained unclassified. For Figure 3 we added the 16 full-length ORF2-intact loci to the full-length ORF-disrupted class (73 loci) and the other 9 to the 5′-truncated class (297 loci).

### HERVK (HML-2) allele states

We resolved HERVK allele states with the HERVK classifier of GraffiTE v1.1, which runs under --human. We counted alleles in proviral units: null (no element), solo (solo LTR, zero units), provirus (one LTR-internal-LTR unit) and prov_xN (N tandem units sharing an LTR at each junction). The classifier ran on all 140 LTR/ERVK discovery candidates (|SVLEN| up to 25 kb), including those --human removed. For each candidate, GraffiTE read the element architecture from the raw RepeatMasker output and determined the reference state by masking 1,500 bp of CHM13 either side of the footprint, increasing the windows where the reference element reached the edge. GraffiTE assigned proviral copy-number states first, when |SVLEN| matched a whole number of the reference element’s LTR-to-LTR period (within 50 bp or 0.5% of |SVLEN|, whichever is larger) and one allele carried two or more units. Otherwise, GraffiTE assigned the alleles from the structure of the element in the variant sequence or, when that structure fitted more than one pair of alleles, from the element that CHM13 carries at the site. We labelled each locus by the first property that applied: MEI when a null allele segregated (21 loci), proviral copy number when an allele carried two or more units (3), and dimorphic when a solo LTR and a provirus segregated (7). GraffiTE rebuilt six loci from 14 discovery calls (four from two, two from three) and wrote each locus as one multi-allelic record; the 31 loci span 36 records of the genotyped call set. GraffiTE writes graph genotypes for the loci and withholds them at proviral copy-number loci, where reads from the reference copy also align to the alternative path. Outside the pipeline, we took the genotypes of those three loci from the assembly-based discovery VCF of the same run (PAV). We never combine the two sources at a locus.

### Comparison of HERVK loci with published catalogues

We summed allele counts over the 27 population sheets of Dataset S2 of Wildschutte et al. 2016 (41 HML-2 loci, 13 present in hg19 and 28 absent) and converted the coordinates to CHM13.0 with UCSC liftOver (Jun 27, 2025) and the hg19ToHs1 chain; liftOver converted 38 of the 41. We matched converted loci within 1 kb of our locus footprints (the union of member-record spans) with bedtools v2.31.1. For the three unconverted loci, we aligned 5 kb of hg19 sequence flanking each side to CHM13.0 with blastn 2.16.0, mapped 400 kb of hg19 sequence around HERVK115 with minimap2 (2.30-r1287) -x asm5 to exclude the 8p23.1 paralogues, and aligned our inserted sequence to the hg19 K115 provirus. These alignments placed two of the three loci. We compared element form (solo LTR or two-LTR provirus) with their 31 converted non-reference insertions within 100 bp, and took two-LTR carrier frequencies from their Dataset S3. We attached a published name when the converted hg19 coordinate of an HML-2 locus from Subramanian et al. 2011, or of a Neanderthal or Denisovan locus from Agoni et al. 2012 and Marchi et al. 2014, fell within 2 kb of a footprint. Wildschutte et al. report the allele that is non-reference in hg19, so we expressed our frequencies in that direction: insertion frequency where hg19 lacks the element, empty-site frequency where hg19 carries it. We computed Pearson and Spearman correlations over the 17 MEI loci matched to Dataset S2 in R 4.3.1.

### Genomic distribution of MEIs

We partitioned chr1-22, chrX and chrY of CHM13.0 (3,117,275,501 bp) into non-overlapping categories built from the RefSeq Liftoff annotation of CHM13.0 (v5.2) and MANE Select v1.5 (Morales et al. 2022). We matched MANE accessions to primary Liftoff gene copies and recovered 19,086 of 19,293 transcripts. The categories were CDS, 5′UTR and 3′UTR (exonic sequence outside the CDS, assigned per transcript and by strand), a residual exonic class, promoter (the 1 kb immediately upstream of the MANE TSS), intron, non-coding genes and intergenic sequence. We resolved overlaps in that order of priority. We placed each MEI at its VCF breakpoint (POS) and assigned it to the category containing that base. We excluded the 87 SVA VNTR-only records and all 31 HERVK loci, whose 21 MEIs are too few to test, leaving 5,724 MEIs (4,908 Alu, 704 LINE1 and 112 SVA). We also excluded the 487 MEIs whose breakpoints fell in centromeric satellite arrays (CHM13 cenSat track) and tested the remaining 5,237 (4,521 Alu, 610 LINE1 and 106 SVA). For each family and for all MEIs pooled, we built a null distribution by shuffling the breakpoints of each chromosome 1,000 times by one random offset within its sequence outside cenSat, following the circular randomization of regioneR (Gel et al. 2016), which preserves the number of variants per chromosome and their relative spacing. We built a second null of 1,000 permutations from random positions outside cenSat on the same chromosome with the same GC content (1% bins) in the 1 kb flanking the breakpoint (for deletions, flanking the deleted element). We report log2((observed + 0.5)/(mean permuted + 0.5)) and a two-sided empirical P value equal to twice the smaller tail proportion, each tail computed as (number of permutations at or beyond the observed count + 1)/1,001. We corrected P values for the 32 tests of each null (four variant sets by eight categories) with the Benjamini–Hochberg procedure.

We obtained the ENCODE V4 registry of candidate cis-regulatory elements (cCREs; 2,348,854 elements on GRCh38, file ENCFF420VPZ) and lifted it to CHM13 with the UCSC hg38ToHs1 chain and rtracklayer (v. 1.60.1), keeping the elements whose lifted pieces lay on one chromosome and covered at least 95% of their original length (2,341,420 elements, 99.7%). We considered a cCRE active in blood when at least one of 54 biosample-specific cCRE sets from untreated peripheral blood leukocytes (neutrophils, CD14+ monocytes, B cells, natural killer cells, CD4+ and CD8+ αβ T cells, and peripheral blood mononuclear cells) reported a high signal for it (DNase, or H3K4me3, H3K27ac or CTCF where DNase was not assayed); 612,727 cCREs passed this rule. Each base of CHM13 was then assigned one category: one of the eight registry classes (restricted to blood-active cCREs in the blood set), a cCRE not active in blood (blood set only), or no cCRE. As for the genomic compartments, each MEI was placed at its breakpoint (VCF POS). We excluded the cenSat regions and the CHM13 intervals of at least 10 kb with no aligned hg38 base (456.5 Mb in total), together with the 488 of the 5,724 Alu, LINE1 and SVA MEIs whose breakpoints fell in them. For the remaining 5,236 MEIs, we compared the observed counts with the circular and GC-matched nulls described above, restricted to the unmasked sequence. We ran the test on insertions and deletions together and separately, computed log2 fold enrichment and two-sided empirical P values as above, and corrected the P values with the Benjamini–Hochberg procedure within each combination of cCRE set, null and variant type.

### Insertion frequency spectrum and discovery curves

We computed each MEI’s frequency from the graph genotypes as the number of TE-bearing alleles over the number of called alleles at that locus, so the denominator falls below 40 at loci with missing calls and on chrX. Because human MEIs are almost free of homoplasy, we treated TE presence as the derived state: the frequency is the alternate allele frequency for insertions and one minus it for deletions relative to CHM13.0. For HERVK we counted solo-LTR and proviral alleles as TE-present and kept only the 21 loci that segregate an empty allele, leaving out 7 dimorphic and 3 copy-number loci. Of the 5,745 MEIs, we dropped 34 at which no genotyped haplotype carried the TE (27 Alu and 7 LINE1), leaving 5,711, and plotted the spectrum in 39 equal-width frequency bins, split by SV type. For the discovery curves we counted, for each family, the MEIs carried by k haplotypes, and passed these histograms to panacus growth v0.2.5 (Parmigiani et al. 2024) with a coverage threshold of 1 and no quorum. Panacus computes, for each m, the expected number of MEIs present in a random subset of m haplotypes. As for the CpG panepigenome, we fitted Heaps’ law with a quadratic term, log V(m) = log K + γ log m + δ (log m)², to each curve by least squares.

#### CaG-private MEIs

To identify MEIs not found in HPRC release 2, we compared the GraffiTE MEIs with SVs from a minigraph-cactus graph (Hickey et al. 2023) built from the HPRC release 2 assemblies (232 individuals) together with the 20 CaG genomes. We used cactus-graphmap, cactus-graphmap-split, cactus-align, cactus-graphmap-join, to obtain a joint GFA and its decomposition to a VCF. We popped bubbles with vcfbub to keep only top level variants. From that VCF we kept SVs with a reference or alternate allele of at least 50 bp that no HPRC haplotype carries, removed those that overlap an SV found in HPRC, which left 5,117 CaG-unique SVs. We matched the 5,842 GraffiTE records against these SVs with truvari bench v5.4.0 (English et al. 2022), which kept the 4,944 SVs of at least 50 bp. We allowed up to 500 bp between breakpoints and required at least 50% sequence and size similarity, with no minimum reciprocal overlap, no maximum size and one-to-one matching (--pick single). We called a GraffiTE record CaG-private when it matched a CaG-unique SV.

### Dating of Segregating TE insertions

#### Ancestral recombination graphs (ARG, SINGER)

We dated each MEI by placing it on a branch of its local genealogy (Table S5). Carriers were the haplotypes carrying the TE (the alternate allele for insertions, the reference allele for deletions); we phased heterozygous graph genotypes with the PAV call at the same locus. For each locus we took the SNVs of the 40 haplotypes in a 2-Mb window centred on the breakpoint, removed all TE records, and sampled ancestral recombination graphs with SINGER v0.1.9 (Deng et al. 2025), setting Ne = 20,000, μ = 1.25 × 10⁻⁸ per site per generation and the window’s mean recombination rate from a CHM13.0-lifted genetic map. We ran 25 MCMC samples with a thinning interval of 20 and kept the last 20 genealogies. On each genealogy we gave each branch e a posterior weight proportional to length(e) × ε^d(e) × (1 − ε)^(n − d(e)), with ε = 0.02, n the number of called haplotypes and d(e) the number whose carrier status disagrees with the leaves below e; the age was uniform along the branch. We averaged the mixtures over genealogies and reported the posterior mean (29 years per generation) and central 95% credible interval.

### Recombination clock (GEVA)

We also estimated allele ages with GEVA (Albers and McVean 2020), using the authors’ program (commit edeafb7) compiled from an unmodified source. For each chromosome we built a phased VCF of the 40 haplotypes from the PAV SNV calls and wrote missing or non-callable genotypes as reference, since GEVA has no missing-data model. We added each TE record as a biallelic site with the TE-bearing allele as the derived (alternate) allele, swapping the alleles of deletion records, and set heterozygous carriers that we could not phase to homozygous reference. We took genetic distances from the Beagle GRCh38 map, lifted to CHM13.0 with the UCSC hg38ToHs1 chain, keeping the longest run of markers with non-decreasing genetic position. We ran GEVA with --Ne 20000, the value implied by the cohort’s nucleotide diversity (θ = 4Neμ ≈ 1 × 10⁻³), --mut 1.25e-8, the distributed HMM files and the default limits of 100 concordant and 100 discordant pairs. We report the posterior mode of the joint clock on filtered pairs, at 29 years per generation.

### Molecular clock

Additionally, we dated MEIs from linked SNV divergence on carrier haplotypes, adapting Baduel et al. 2021 to phased, haplotype-resolved human data. Because calls were polarised against CHM13, the TE-bearing allele was the alternate allele for insertions (SVTYPE=INS) and the reference allele for deletions (SVTYPE=DEL); carrier haplotypes were those carrying the TE-bearing allele, and records with fewer than two carriers were left undated. We phased heterozygous graph genotypes against the PAV calls of each haplotype (breakpoints within 200 bp, SVLEN within 30%); about 0.9% of carrier haplotypes could not be resolved, and we dropped them. For each pair of carrier haplotypes we counted differing biallelic SNV sites within a 70-kb window centred on the breakpoint (excluding 50 bp either side of the breakpoint, and for deletions the deleted element) and divided by the jointly callable length *L*, defined as the window intersected with the PAV callable regions of both haplotypes minus the excluded sequence. Pairs with <10 kb of jointly callable sequence were excluded, and insertions with no qualifying pair were undated. Per-insertion divergence was summarised as the maximum pairwise per-site divergence across carrier pairs, *d*max, representing the deepest carrier coalescence and thus a lower bound on insertion age. Age in generations was computed as *T* = *d*max/(*k*μ) with *k* = 2 (accounting for independent substitution along both lineages) and μ = 1.25 × 10⁻⁸ substitutions per site per generation (pedigree rate; Kong et al. 2012), converted to years assuming a 29-year generation interval (Fenner 2005); ages scale linearly in μ. A two-sided 95% exact Poisson interval (Garwood 1936) on the pairwise difference count for the *d*max pair was propagated through *L* and the age equation, capturing finite-site sampling uncertainty only.

### Panepigenome analysis of CpGs in TEs

To compute a panepigenome growth curve for each TE family, we first calculated the frequency histogram of each CpG in each family across all copies that satisfy our filter. Then, we used `panacus growth` (Parmigiani et al. 2024) to generate the exact growth curve. To fit Heap’s law with quadratic correction (Fontanelli and Li 2025), we used the linearized model and estimated the parameters with `lm()` from the growth curves generated by panacus for each TE family: *log(V(n)) = log(K) + γ · log(n) + δ · log(n)²* where *V(n)* is the number of CpGs discovered after *n* genomes. We interpret the effective *γ* (*γ__eff_*) and *δ* at the last observed genome, where *γ__eff_(n) = γ +* 2*δ · log(n).* If *δ >* 0, the pangenome grows faster than Heaps’ Law. When *δ <* 0, it grows slower than Heaps’ Law.

### Epigenetic mapping to TEs with GraffiTE

To map methylation to TE polymorphisms, we employed GraffiTE’s new feature. That is, GraffiTE now realigns the long-reads to the graph genome with `vg giraffe` (Sirén Jouni et al. 2021), computes methylation summary statistics over the graph and projects these to the polymorphic TEs in each sample with `panmethyl` (Groza et al. 2025). The results is a VCF and BED describing the methylation levels of each TE in each sample:

`nextflow run cgroza/GraffiTE --genotype_with methylation_bams.csv`

`--epigenomes --graffite_vcf pangenome.vcf --reference CHM13.fa`

`--genotype true --graph_method giraffe --code C+m. --bed CHM13_repeats.bed`

`To generate Bandage illustrations of TE insertions, we used chop_gfa.py and translate_lifted.py.`

To compare GraffiTE estimates to pb-cpg-tools, we mapped the reads to CHM13, annotated methylation levels with `aligned_bam_to_cpg_scores --pileup-mode count --modsites-mode denovo,` and computed the mean methylation level over repeats with `bedtools map`. Here, we used the RepeatMasker annotation of CHM13, and selected only Alu, LINE1, SVA, and ERVK repeats. Hypomethylated TEs are defined as the 10th percentile within each family. We tested differences in the mean methylation level between polymorphic and fixed TEs using the t-test and corrected p-values with Benjamini-Hochberg (Benjamini and Hochberg 1995). For the above analyses, all TEs were filtered for a minimum 250 bp in length and an average read depth of 4 reads over the CpGs. We defined the methylation rate as the methylation count of a CpG divided by its allele count. The methylation count of a CpG is the number of its epialleles with a methylation level above 50%.

### Analysis of TE age with respect to methylation

To infer the effect of TE age on its methylation level, we used beta regression with the `betareg` R package (Cribari-Neto and Zeileis 2010). The use of beta regression is appropriate because the methylation levels are between 0 and 1 and the beta distribution fits the observed distribution of TE methylation levels. We employed the following model that includes TE age in log10 years and corrects for TE family, donor age, the local recombination rate measured over 500 kbp, GC content, genomic context and the branch posterior probability from the ARG model (the highest posterior probability of the TE’s branch across the 20 genealogies). We also corrected for donor blood composition:

betareg(data, formula = TE_methylation ∼ log10(Age_years) + TE_family + Donor_age + Recombination_rate + GC_content + Genomic_context + ARG_posterior_probability + EOS_p + BASO_p + NEU_p + LYM_p + MONO_p).

We fitted the model to all dated MEIs, once with each set of ages (ARG, molecular clock and GEVA). With the molecular clock, we removed TEs with an estimated age of 0, at which no SNV differs between carrier haplotypes. To interpret the effect size of age on methylation, we calculated the expected methylation levels at age 0 and after 1 Mya for each family.

### Carrier fraction

For each MEI we computed the carrier fraction as the number of phase-resolved haplotypes carrying the TE-bearing allele divided by the number of haplotypes at the locus (40 on autosomes; 30 on chrX, where the 10 male genomes contribute one haplotype each).

### LINE1 activity potential

We labelled each of the 704 LINE1 MEIs by its sequence structure (see LINE1 insertions). We labelled an element theoretically active when it was full-length intact: it started within the first 50 bp of the L1.3 consensus, covered at least 95% of it, carried no 5′ inversion, and had intact ORF1 (338 amino acids) and ORF2 (1,275 amino acids), each with a single peptide covering at least 90% of the reference protein, anchored at both termini, at 95% or higher amino-acid identity.

#### Use of generative AI

We used Claude Code (Anthropic) with the models Claude Opus 5 and Claude Opus 5.5 to (i) write and debug parts of the code of GraffiTE v1.1 and of the analyses (call-set curation, TE dating, TSD and HERVK analyses, figures 2, 3 and 5B), (ii) run these analyses on our data under our direction, and (iii) draft and edit related parts of the Methods, Results and figure legends. We reviewed the code and text it produced, checked its results against the underlying data, and take full responsibility for the content of this article.

## Code and data availability

GraffiTE v1.1 is available on GitHub (https://github.com/cgroza/GraffiTE); this paper describes release v1.1.0.0. The headers of the released VCFs name the development versions our analyses used. The scripts that build the curated call set and every figure are in the paper/CaG2026 directory of the same repository. CARTaGENE long-reads, methylomes, assemblies and and per sample genotypes and derived tables can be requested at https://cartagene.qc.ca.

## Contributions

C. Groza designed and implemented the epigenetic pipeline and related results. C. Goubert designed and implemented the annotation and dating pipeline and related results. Both C. Groza and C. Goubert conceptualized analyses, created figures and drafted the manuscript. R. Laflamme and G. Lettre designed and implemented the sample selection and contributed to variant annotation. G. Bourque, G. Lettre and J. Ragoussis initiated, designed and directed the sequencing of the CARTaGENE samples used in this study. R. Eveleigh participated in the sequencing of CARTaGENE. T.J. Wheeler supported the repeat annotation work. All authors contributed to study design and manuscript draft.

