## Supplemental figures for "Integrated genotyping, annotation and epigenetic characterization of transposable elements with GraffiTE"

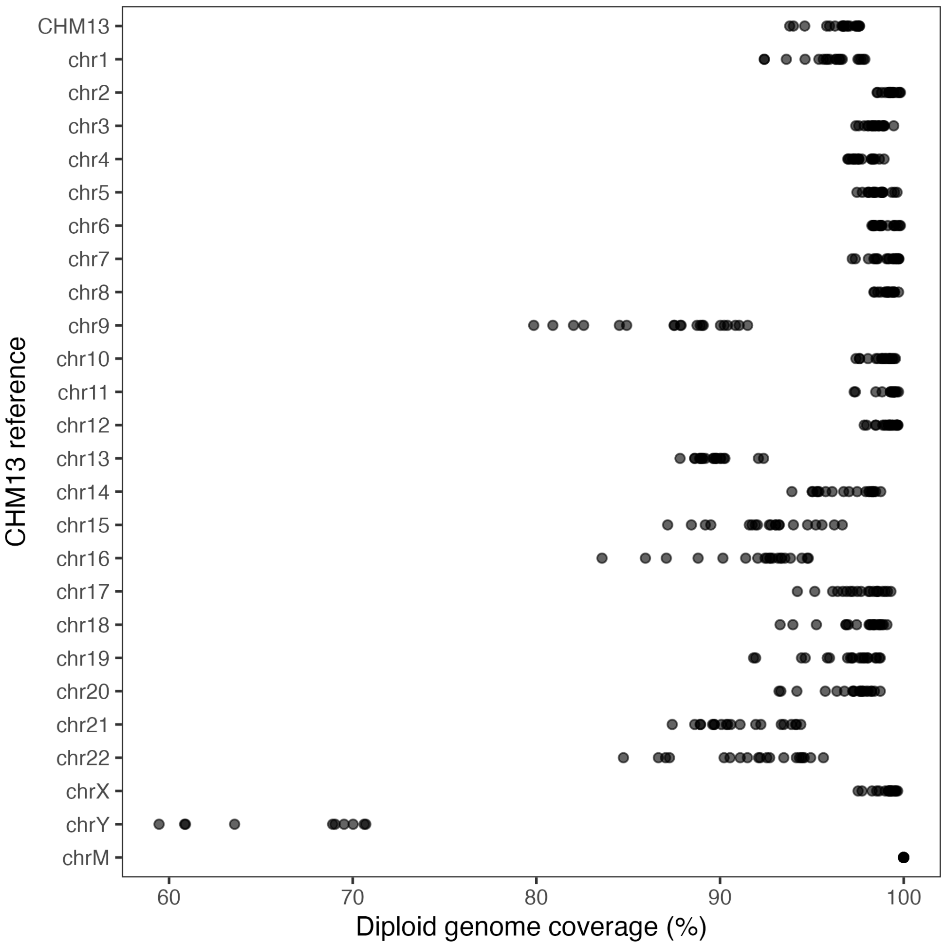


Fig S1: Percent of CHM13 contigs spanned by each assembled diploid CARTaGENE assembly.

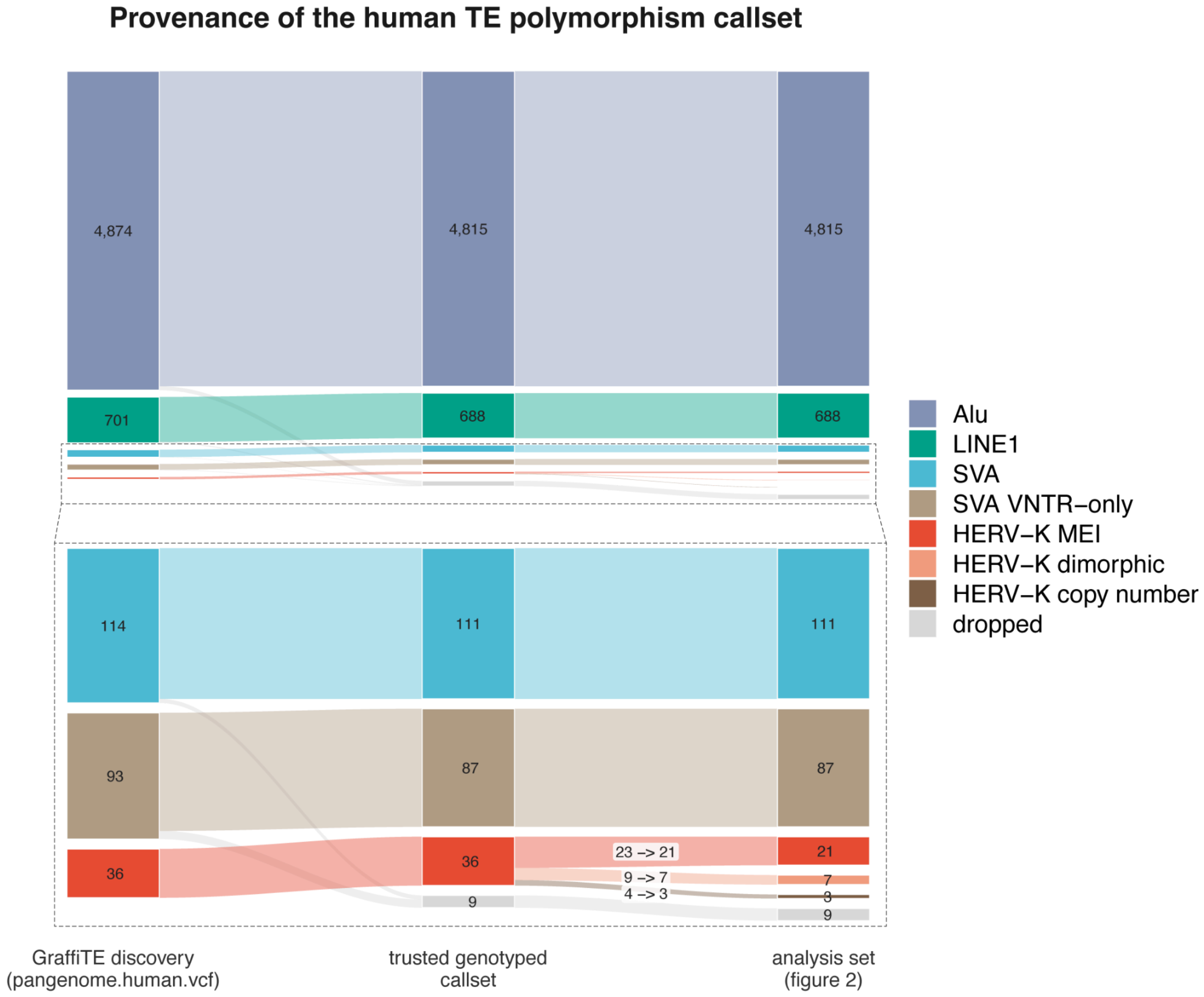


Fig S2: Provenance of the human TE polymorphism callset. Nine records from the human discovery VCF have no vg call record (no graph genotype); no HERV−K record is dropped. The 36 HERV−K records resolve to 31 loci: one reference element reached by two or more graph−derived calls placed at different breakpoints. The 87 SVA VNTR−only records are retained in the callset but excluded from analyses that treat a record as MEI, as are the 10 HERV−K which segregate either as dimorphic (solo-LTR vs proviral) or as proviral CNV.

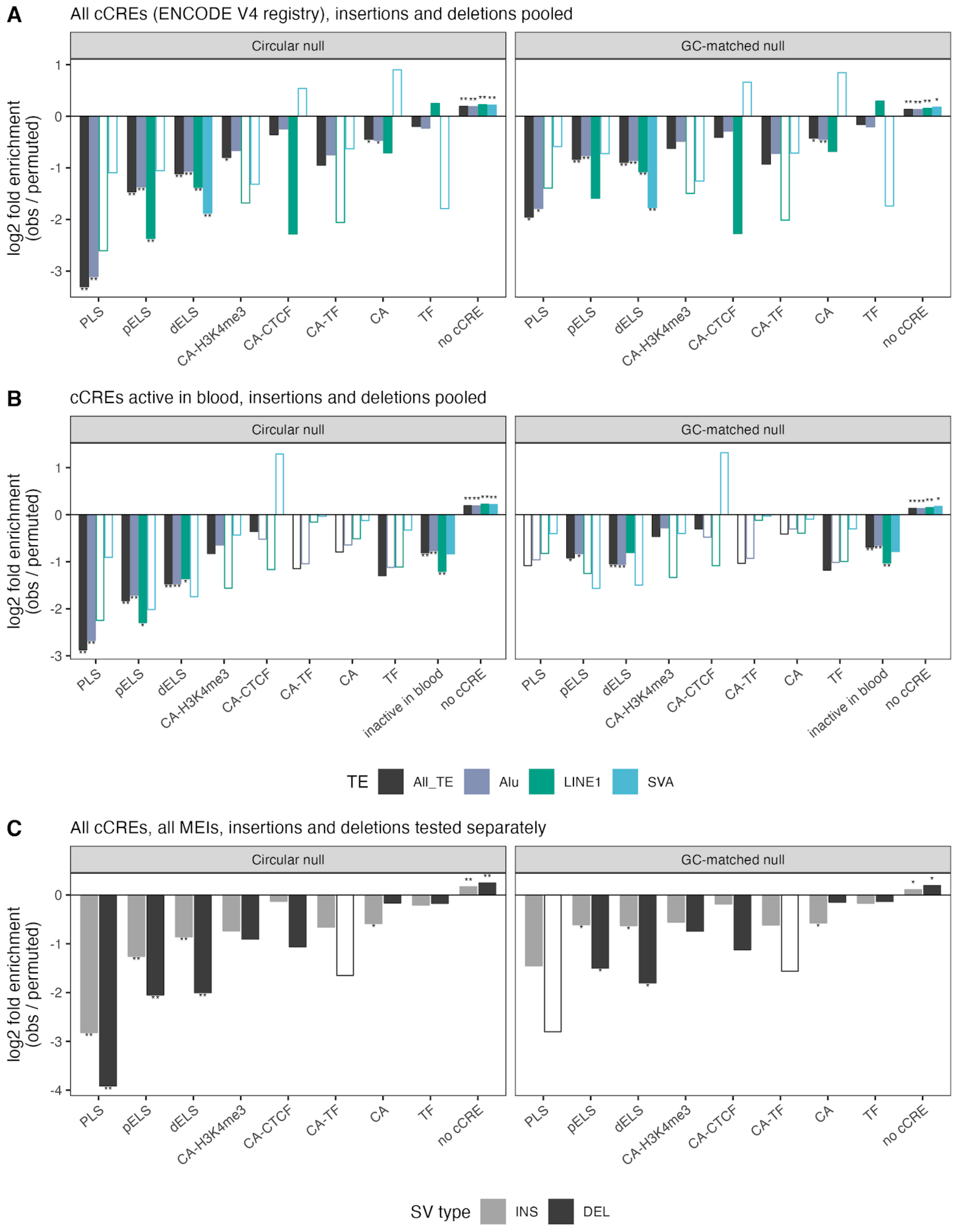


Fig S3: MEI enrichment/depletion in ENCODE candidate cis-regulatory elements (cCREs). Log2 fold enrichment (observed/permuted) of MEI breakpoints in each class of the ENCODE V4

cCRE registry lifted to CHM13: PLS, promoter-like; pELS, proximal enhancer-like; dELS, distal enhancer-like; CA-H3K4me3, CA-CTCF and CA-TF, accessible chromatin with H3K4me3, CTCF or transcription factor binding; CA, accessible chromatin only; TF, transcription factor binding without high accessibility; no cCRE, sequence outside all cCREs. Left: circular null, with the breakpoints of each chromosome shifted by one random distance; right: random positions matched for the GC content of the sequence flanking each breakpoint (Methods). A) All cCREs, insertions and deletions pooled (n = 5,236 MEIs: 4,520 Alu, 610 LINE1 and 106 SVA; All_TE, the three families pooled). B) cCREs active in at least one of 54 ENCODE blood leukocyte samples (Methods); "inactive in blood" groups the other cCREs of the registry. C) All MEIs, with insertions (n = 3,675) and deletions (n = 1,561) tested separately. Open bars: fewer than five breakpoints expected under the null. Breakpoints in cenSat arrays and in CHM13 intervals of at least 10 kb with no hg38 alignment were excluded. *: P-adj (B.H.) < 0.05, **: P-adj < 0.01.

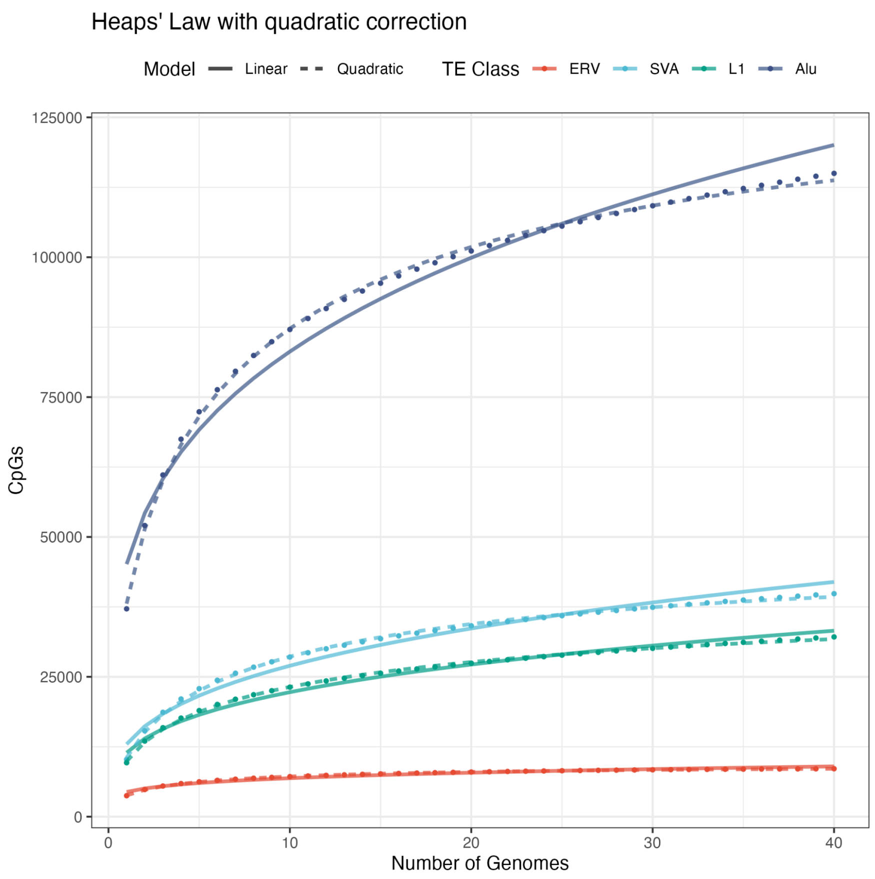


Fig S4: Heaps’ Law fits to the growth curves inferred by Panacus for each TE family (dots). The fits with quadratic correction (dashed lines) are contrasted to uncorrected plain Heaps’ Law (full lines).

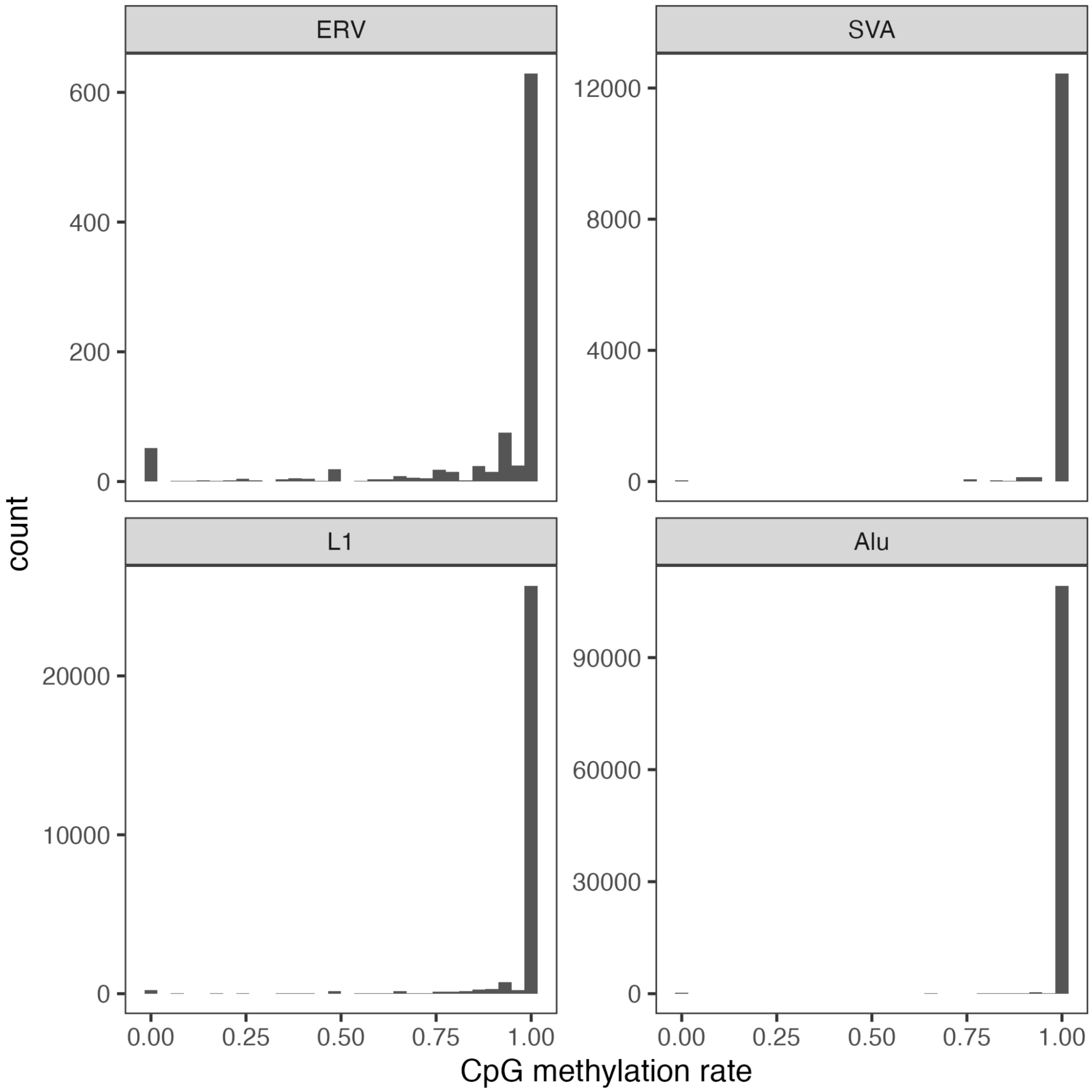


Fig S5: Histograms of methylation rates of each CpG (methylation count divided by allele count) associated with the cumulative distributions shown in Fig 3E.

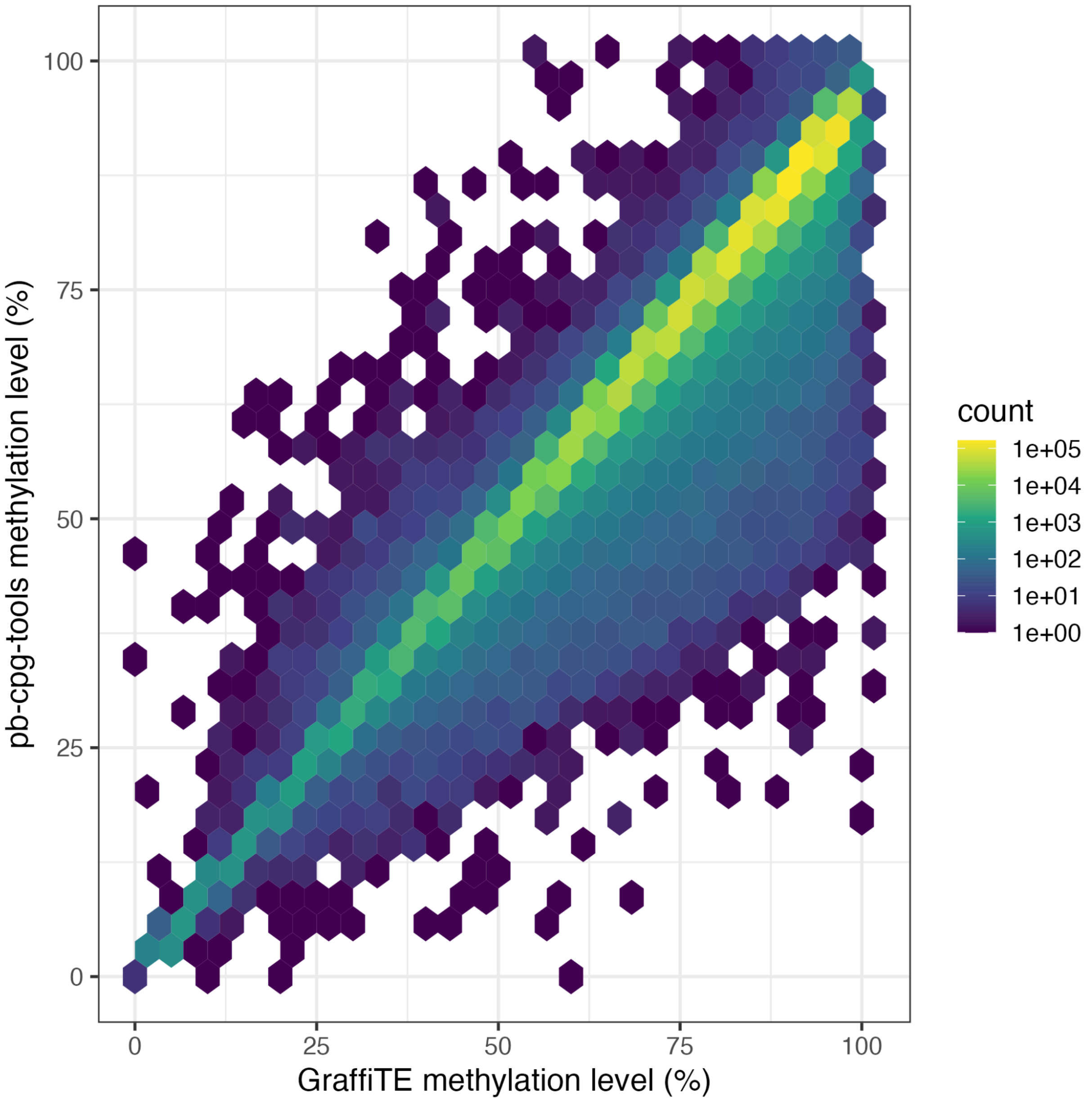


Fig S6: Comparison of average TE methylation levels between `pb-cpg-tools aligned_bam_to_cpg_scores` and GraffiTE over CHM13 TEs annotated with RepeatMasker.

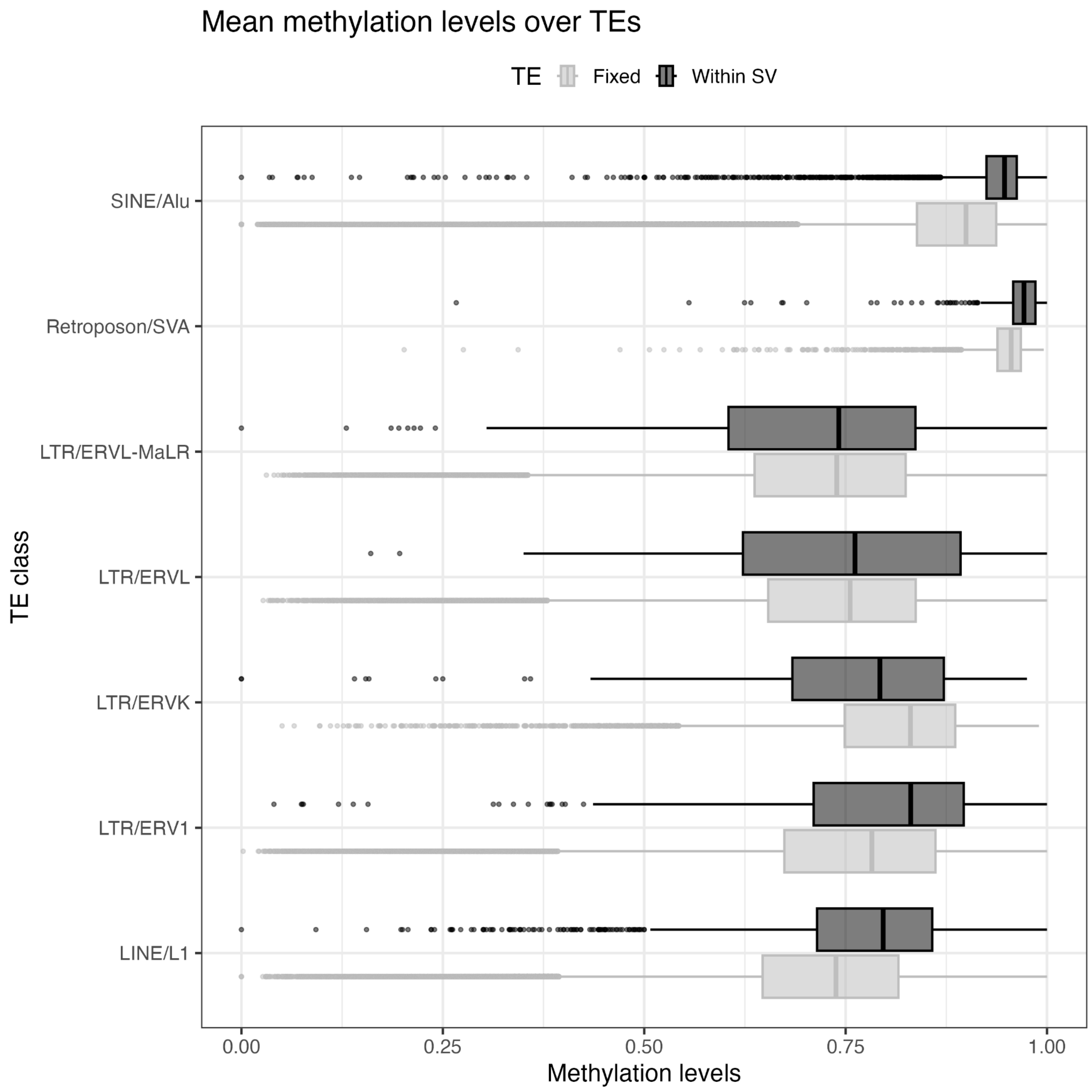


Fig S7: A) Average CpG levels methylation across fixed and variable across all TEs annotated by GraffiTE, including those not in the trusted set. This includes TEs that are part of other SVs and are not the result of transposition activity and also additional families of ERVs.

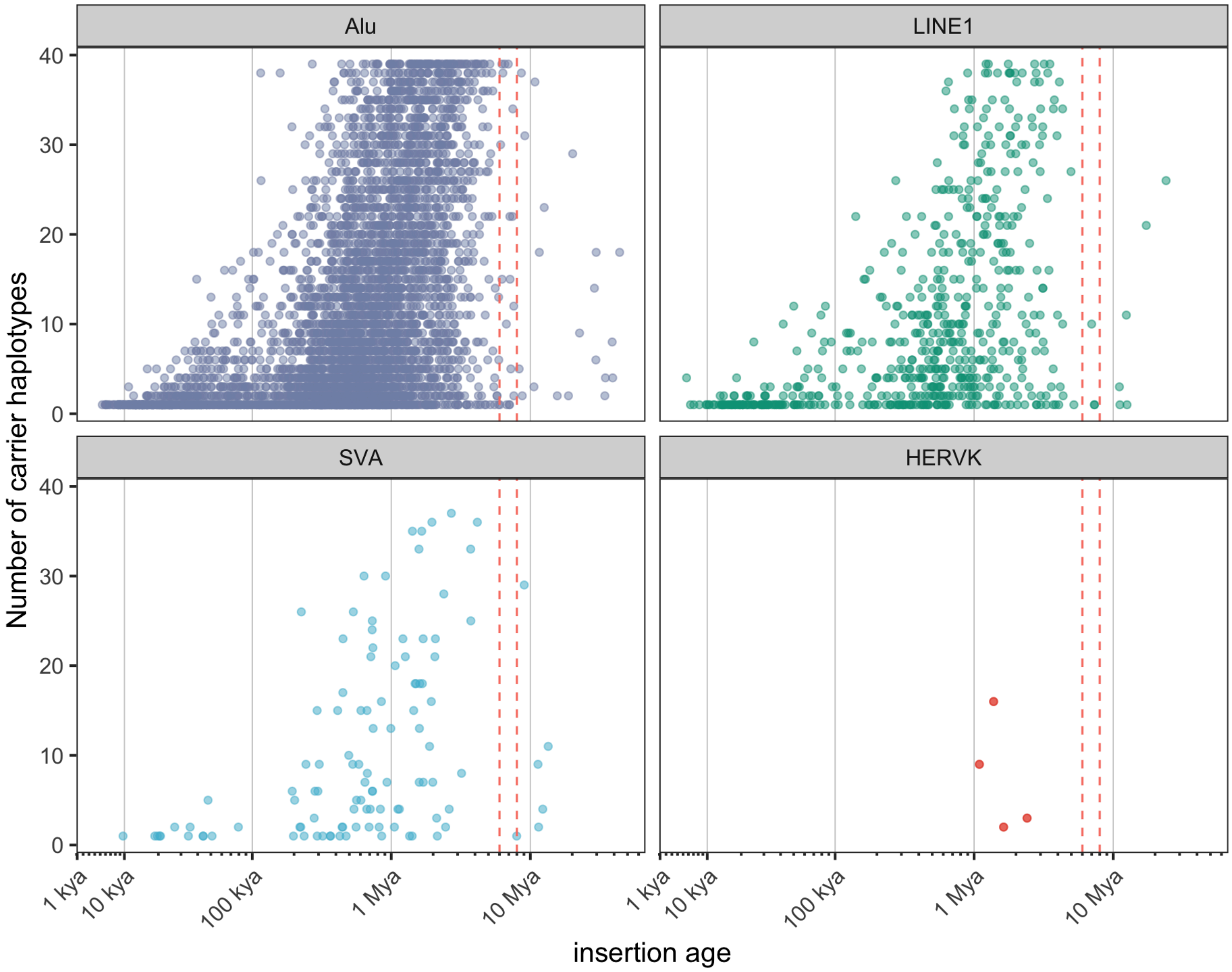


Fig S8: Dated TE ages compared to their allele count in CARTaGENE, stratified by TE family.

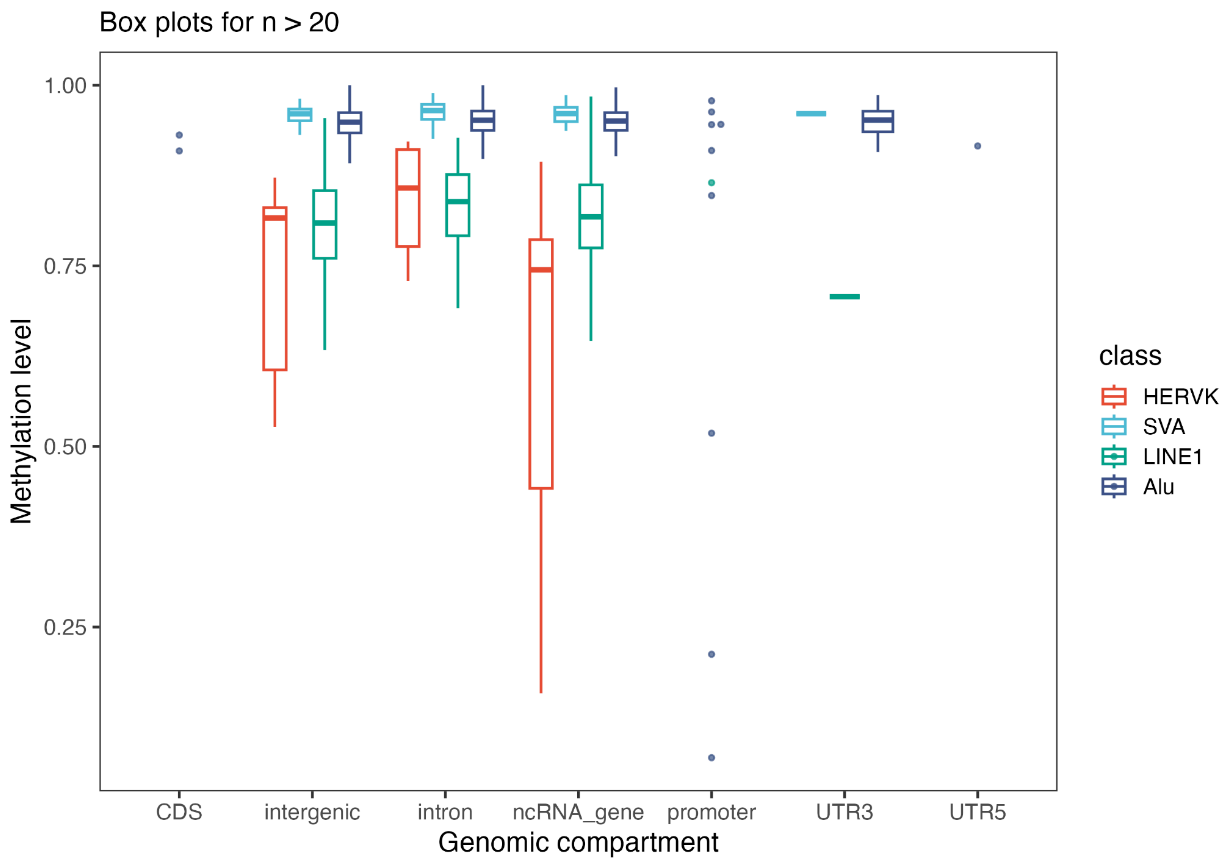


Fig S9: The average methylation levels of each TE across CARTaGENE, stratified by genomic context and family.

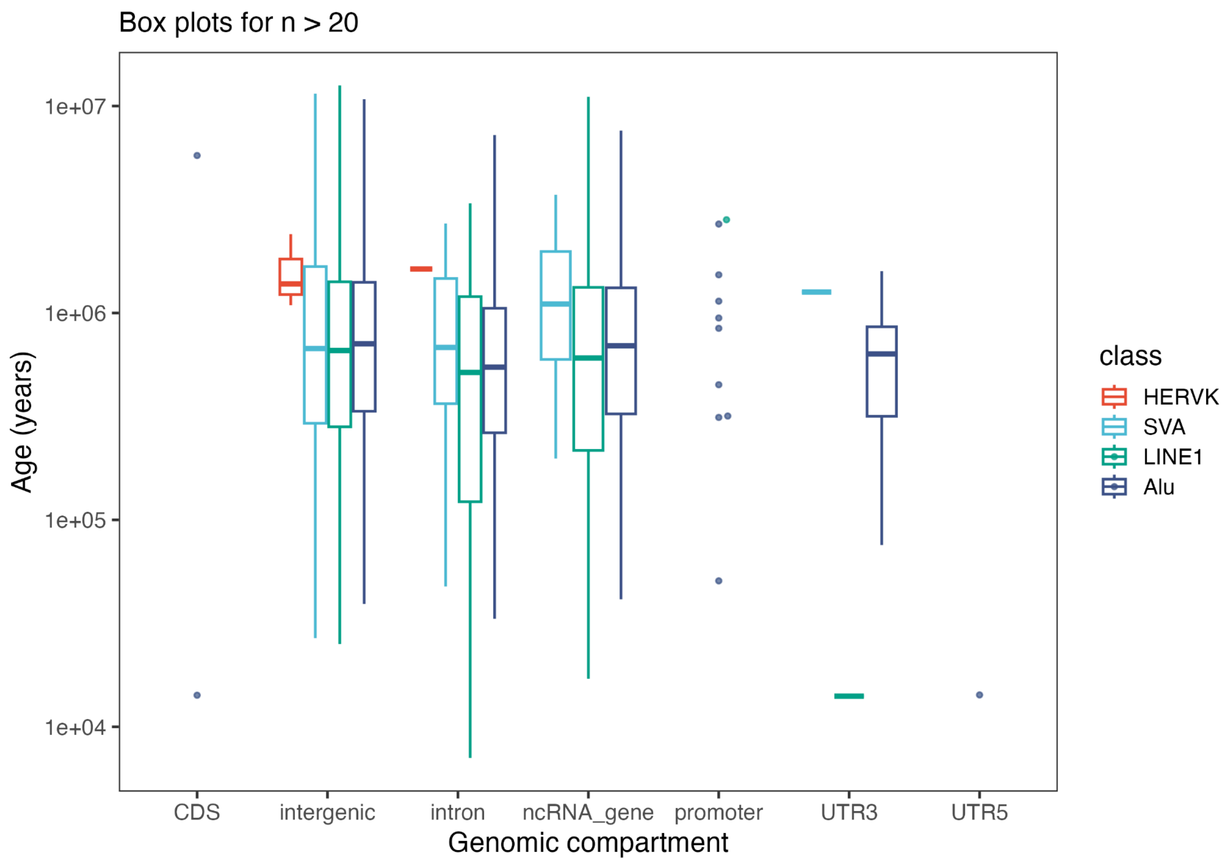


Fig S10: The ages of dated TEs, stratified by genomic context.
